# From prehistory to present-day: How isolation shaped the distinct genomic makeup of Italian Alpine valleys

**DOI:** 10.64898/2026.07.16.738935

**Authors:** Giacomo Villani, Nicola Rambaldi Migliore, Anna Tommasi, Nicola Nannini, Erika Partel, Vittoria Cerizza, Ilaria Piloni, Gary Sorasio, Valeria Nicolini, Irene Cardinali, Rosalinda Di Gerlando, Giulia Bozzari, Elena Raimondi, Donato Riccadonna, Roberta Raffaetà, Heidi Christine Hauffe, Licia Colli, Anna Olivieri, Antonio Salas, Hovirag Lancioni, Alessandro Fedrigotti, Paolo Ajmone Marsan, Antonio Torroni, Alessandro Achilli

## Abstract

Historically, the eastern Italian Alps have provided crucial geographic corridors for cultural and genetic exchange between the Mediterranean region and Central and Northern Europe. Although recent archaeogenomic data suggest a strong regional persistence of Anatolian Neolithic ancestry followed by the arrival of Yamnaya-related components during the Bronze Age, the genomic landscape of modern Alpine populations remains largely unmapped. In this study, we verified the persistence of these signals in the present-day Rendena and Ledro Valleys by integrating novel complete mitochondrial genomes (N=185) and genome-wide SNP data (N=96), the latter combined with a novel Italian genomic dataset (N=139). Our findings show that the Rendena and Ledro populations form a distinct “modern Alpine” genomic group, which retains a significant proportion of early European Neolithic ancestry, aligning with ancient eastern Italian Alpine individuals and modern Sardinians. More recently, geographic isolation and localized genetic drift appear to have shaped the two valleys differently. Demographic reconstructions reveal asynchronous population declines over the past two millennia, followed by a sharp, synchronized bottleneck 200-300 years ago, which coincided with historical plague outbreaks. Remarkably, this structured drift persisted at an extremely fine microgeographic scale within the valleys, resulting in internal genetic subclusters that directly correlate with local topography. This micro-differentiation was likely maintained by steep geographic barriers and/or local endogamous practices. This study ultimately underscores how geographic barriers and isolation can preserve ancient genomic components and shape highly localized genetic structures over centuries even to the modern day.

## Introduction

Anatomically modern humans appeared in Eastern and Western Europe^1,2^ and in Italy^3^ by at least 45,000 years ago (ya). A simplified, yet not exhaustive, pan-European ancestral scenario suggests that before and after the Last Glacial Maximum (25-22,000 ya), migrations and admixture events resulted in the widespread diffusion of the Western Hunter-Gatherer (WHG) ancestry in Western Europe, including Italy^4–6^. Around 10-8,000 ya, the arrival of Neolithic farmers from Anatolia (Anatolia_N) impacted the European gene pool. This major demographic shift is still evident nowadays in regions such as Sardinia^7^. During the Bronze Age, a Yamnaya-related component from the Pontic-Caspian Steppe, with sources related to Eastern Hunter-Gatherers (EHG) and Caucasus Hunter-Gatherers/Iranian Neolithic (CHG/Iran_N), appeared in Europe^7,8^. In addition to this pan-European history, Italy was shaped by region-specific historical processes, such as the Greek colonization in the south, and the expansion and consolidation of the Roman Republic/Empire^9^.

This complex history has shaped modern-day Italy into a country composed of a mosaic of cultural, linguistic and genetic diversity. Genome-wide studies have revealed patterns reflecting this complex history. A clear latitudinal cline is apparent across modern Italy^7,10–12^, with the island of Sardinia forming a distinct genetic group^13^, while mitochondrial DNA (mtDNA) studies have highlighted finer-scale regional heterogeneity^14,15^. However, despite several investigations focusing on regions such as Sicily^14^, Piedmont^16^, Tuscany^17,18^ and Umbria^15^, much of Italy’s micro-geographic genetic structure remains underexplored. Of particular interest are the eastern Alpine valleys in Trentino-Alto Adige/Südtirol (TAA). Located at the interface between the Italian peninsula and Central Europe, this region has long served as a contact zone from the Chalcolithic through the Roman and Medieval periods, it was part of the Holy Roman Empire and was annexed to Italy after the First World War. A recent analysis of ancient genomes from this eastern Alpine region revealed a Mesolithic admixture of WHG and EHG, followed by a period of genetic continuity stretching from the Neolithic to the Bronze Age, interrupted with the arrival of the Yamnaya-related component around 4,400 ya^19^.

These genomic profiles, before the Bronze Age, align with that of the Tyrolean Iceman (hereafter referred to as the Iceman), also known as “Ötzi,” the world’s oldest glacier mummy who lived during the Chalcolithic and exhibited high Anatolia_N-related ancestry and low hunter-gatherer ancestry^19–21^. However, studies of modern populations in the Alpine valleys have never achieved this level of genomic detail.

The Rendena (REN) and Ledro (LED) Valleys provide interesting case studies for investigating fine-scale demographic and genetic patterns. Located approximately 20 kilometers apart, the present-day communities of these valleys share similar features, including relative geographic isolation, strong links to agricultural traditions, and dependence on water resources. They also have some cultural and microgeographic peculiarities. Alongside Italian, the Ledrensi speak a relatively uniform Lombard dialect, while the Rendeneri speak local dialects influenced by Ladin in the north and eastern Lombard dialects in the south. The Rendena Valley runs north-south along the Sarca River (Fig. 1A), while the Ledro Valley runs east-west and consists of an interconnected system of sub-valleys, namely (Ampola, Concei, and Ledro) (Fig. 1A). The discovery of an extensive 4,000-year-old pile-dwelling settlement along Lake Ledro in 1929 highlights its deep prehistoric roots.

**Figure 1.**
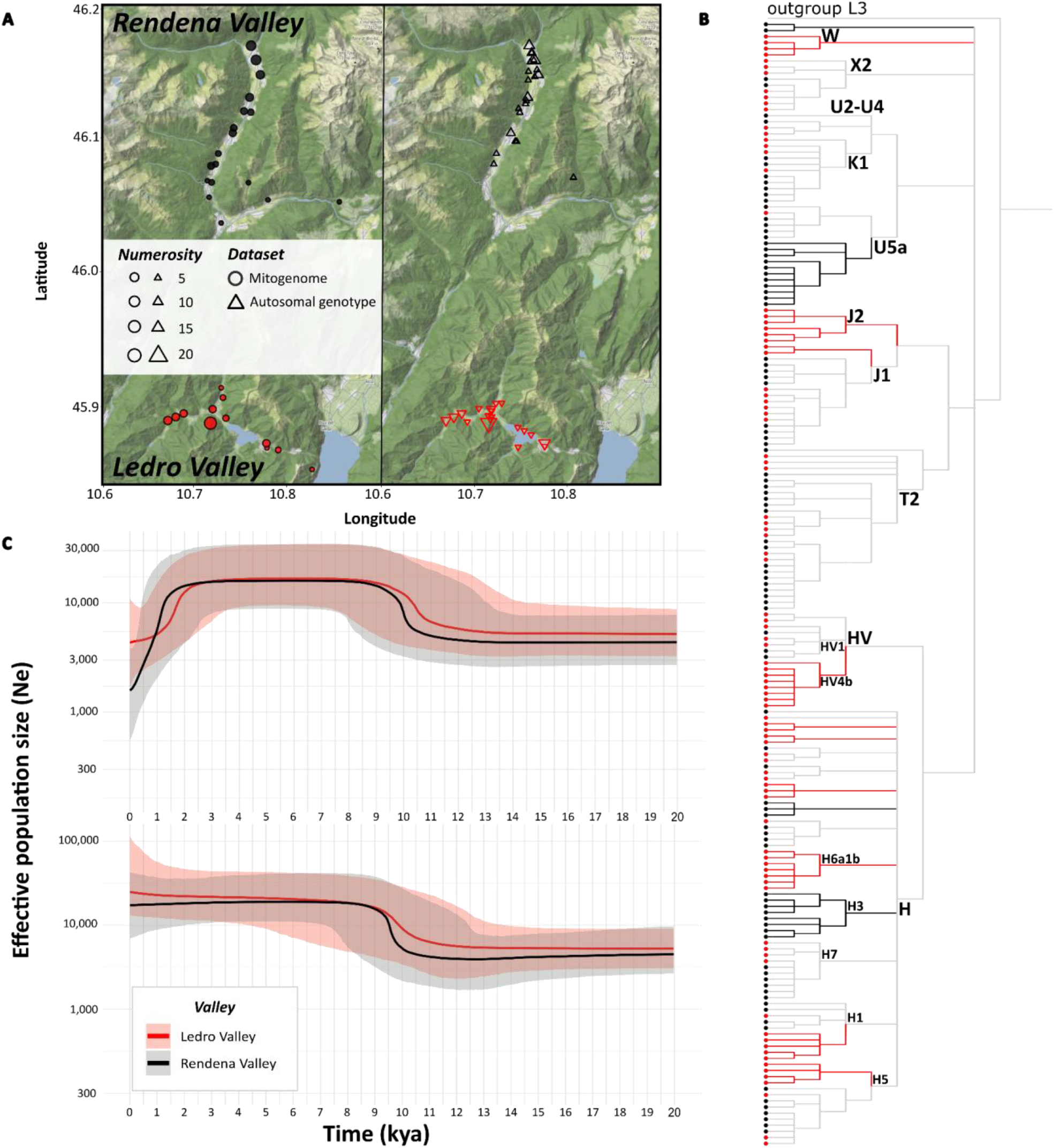
Geographic origin of mitogenomes and autosomal genotypes in the Rendena and Ledro Valleys; mtDNA phylogeny and demographic history of maternal lineages. **A)** Geographic origin of the individuals whose mitogenomes and autosomal genotypes were analyzed in this study. Black and red circles indicate the geographic origin of the terminal maternal ancestor (TMA) of individuals from Rendena Valley (N = 94) and Ledro Valley (N = 91), respectively. Black upward and red downward triangles are positioned at the mean latitude and longitude calculated from grandparents originating from the valley for individuals from Rendena Valley (N = 37) and Ledro Valley (N = 43), respectively. See Dataset S1 for further details. **B)** Maximum parsimony (MP) tree including all mitogenomes from the Rendena (black circles) and Ledro (red circles) Valleys, plus a published Sicilian L3 outgroup. The most frequent haplogroups are labeled along the corresponding branches. Branches corresponding to mitochondrial lineages found exclusively in Rendena Valley are highlighted in black, whereas those found exclusively in Ledro Valley are highlighted in red. See Dataset S2 for additional details. **C)** Bayesian Skyline Plots (BSPs) showing changes in effective population size over time for the two valleys, computed using 1500 time windows. The upper panel shows analysis performed using all mitochondrial genome sequences from each valley (N_REN = 94 and N_LED = 91), whereas the lower panel shows analysis performed by assessing only the unique mitochondrial genome sequences and counting each shared haplotype only once (N_REN = 51 and N_LED = 54).

In this study, we first characterized and compared the complete mitochondrial genome in modern individuals from the two neighboring Alpine valleys. Then, we expanded the analysis to include nuclear genomic variation, comparing it with a novel, genome-wide dataset from Italy. This allowed us to detail the demographic, historic, and geographic factors that shaped their present-day gene pools.

## Results

### MtDNA variation of Rendena and Ledro Valleys

We obtained 185 high-quality mitochondrial genomes with an average depth of coverage of 723X (minimum depth of coverage 46X and maximum depth of coverage 1,984X). Ninety-four were from maternally unrelated individuals whose terminal maternal ancestor (TMA) was from the Rendena Valley, and 91 were from individuals with the TMA born in the Ledro Valley (Fig. 1A, left panel; Dataset S1).

These novel mitogenome datasets were characterized by relatively low nucleotide diversity (π = 0.002 in both valleys) and high haplotype diversity (REN-Hd = 0.975 and LED-Hd = 0.986). This pattern typically indicates a recent, rapid population expansion, or “bottleneck recovery”. Among the 94 REN mitogenomes, 51 distinct haplotypes were identified and classified into 35 haplogroups; 62 LED haplotypes were identified from 91 mtDNA sequences and assigned to 41 haplogroups. Four new sub-haplogroups were identified, two of which were shared (J1c3e2a and HV1e), and two were valley-specific (REN: H3ap1; LED: H5t1). All lineages are of Western Eurasian origin^13,15^ and were subsequently grouped into 18 macro-haplogroups based on their phylogenetic relationships and observed frequencies (Fig. 1B, Dataset S2, and Materials and Methods section for further details). In both valleys, the most frequent mtDNA haplogroup is H with its subclades (37% in LED; 41% in REN), followed by haplogroup T2 (18% in LED; 9.8% in REN).

The two valleys have similar frequencies of most macro-haplogroups (13 out of 18), but also some valley-specific lineages, including H3 in REN, and H6a1b, HV4b, J2, and W restricted to LED (Fig. 1B). This mitogenome pattern is also evident in a principal component analysis (PCA) of Western Eurasia and Northern Africa (Figure S1, Dataset S3). Although REN and LED are separate from each other, both stand out from the broader northern Italian mtDNA variation along the first and second principal components.

### Long-term demographic patterns based on mitogenomes

We examined demographic expansion dynamics through Bayesian Skyline Plot (BSP) reconstructions. Both valleys showed a similar trend: a period of relative stability until approximately 10,000 years ago, when the “Neolithic revolution” triggered a marked demographic expansion in just one millennium. Following a second plateau lasting another 8,000 years, a sharp decline was observed from 2,000 ya to present, initially in Ledro and later in Rendena, where it was more marked (Fig. 1C; upper panel). When the BSP analysis was repeated with unique haplotypes only, the recent decline in effective population size was notably absent in both valleys (Fig. 1C; lower panel). This contrasting result indicates that the BSP signal in Fig. 1C (upper panel) was primarily driven by the high redundancy of a subset of mitochondrial haplotypes rather than a true loss of population size, supporting the local isolation hypothesis. Over the past two thousand years, the maternal gene pool has been dominated by lineages that currently exhibit the highest haplotype redundancy (T2, J1, and U5a), and there has been limited gene flow from outside the valleys. Recent isolation has also led to micro-differentiation of the aforementioned valley-specific sub-haplogroups in terminal branches.

### Genome-wide variation of Rendena and Ledro Valleys

Given the current peculiarities in the mitochondrial gene pool of the two valleys with respect to other Italian groups, we selected 48 individuals from each valley, with maximally divergent mtDNA lineages and wide geographical distribution, and an additional 139 individuals from across Italy to further explore their genomic diversity through genome-wide genotyping with the Human Origins SNP Array (HO chip: 629,443 SNPs; Datasets S1 and S4). Individuals were classified according to their genealogical background, particularly looking at the birthplace of the grandparents (GP). Particular emphasis was placed on those with all four GP from the same valley, defined as REN_4GP (N = 26) and LED_4GP (N = 34). All other individuals (i.e., with at least one grandparent from the valley) were classified as REN_XGP (N = 11) and LED_XGP (N = 9), with X indicating the number of grandparents from the valley. The other Italian individuals were selected with all four GP from the same region and at least three GPs born within the same province. All Italian individuals not belonging to the two Alpine valleys were grouped into four macro areas: North, Center, South, and Sardinia, as in^11^.

### Genomic distinctiveness of the two Alpine valleys

We performed an initial exploratory analysis of Western Eurasia through a PCA using available data (AADRv62)^22,23^ and our novel dataset (Fig. 2A). This genetic landscape confirmed that Italian individuals are stretched along a latitudinal cline, with northern Italians aligning more closely with north-western Europeans, and southern Italians with eastern Mediterranean populations, as already observed with a different SNP panel^7,11^ and complete genomes^12^. Instead, LED and REN individuals formed a distinct and compact cluster close to, but not overlapping with, other northern Italian individuals. Interestingly, this Alpine cluster is located between the main Italian clusters and the Sardinian one, suggesting peculiar intermediate genomic features. Modern Sardinians have the highest Anatolian_N ancestry in Western Eurasia and a negligible Yamnaya-related component, likely due to their isolation until the Iron Age. More recent admixture with Mediterranean populations, including Phoenicians and Corsicans, has contributed to their genetic makeup^24,25^.

**Fig. 2.**
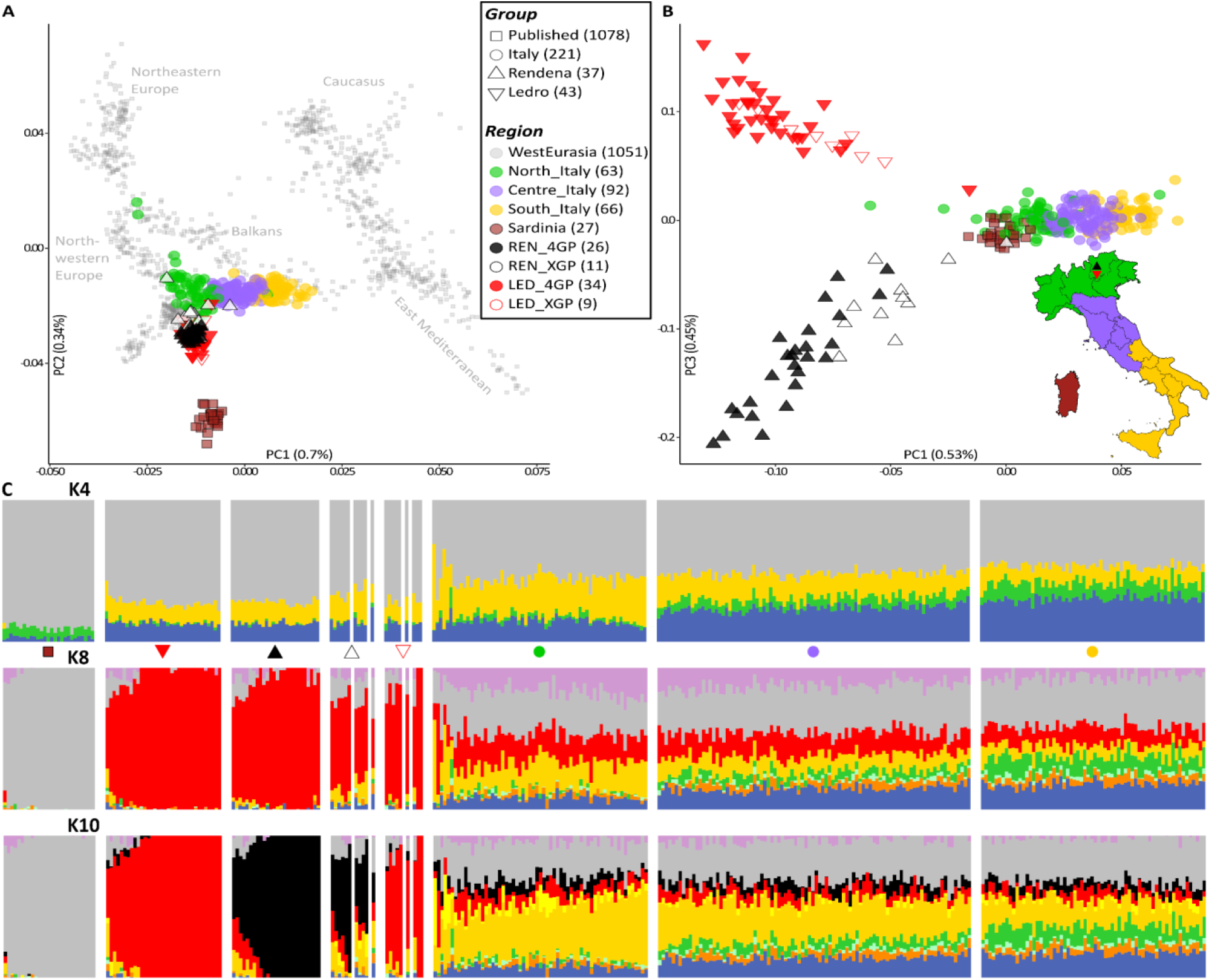
PCA and Admixture analyses. **A)** PCA of Western Eurasia, PC1 and PC2. See also the inset in B for the subdivision of Italian individuals into North, Centre and South Italy. REN and LED individuals were subdivided according to their genealogies, as detailed in the text. **B)** PCA of Italy for PC1 and PC3. Same legend as in A. **C)** ADMIXTURE bar plots of Western Eurasia showing only Italian populations at K=4, K=8 and K=10, same legend as in A. LED_XGP and REN_XGP are ordered from left to right: 3,2,1 GPs born in a specific valley. See Figure S4 for CV error barplot and complete Western Eurasian ADMIXTURE.

When the analysis is restricted to Italy (Fig. S3), the PCA shows that REN and LED are clearly distinct from the rest of Italy and Sardinia. In addition, PC3 further separates the Ledro Valley from the Rendena Valley (Fig. 2B). Considering the genealogical subgroups, LED_4GP and REN_4GP have internal axes of variability, while LED_XGP and REN_XGP are drawn toward the rest of Italy according to their respective genealogies: their grandparents originating outside the valleys are mainly from other valleys in TAA, neighboring regions (Lombardy, Veneto and Friuli Venezia Giulia) and then other Italians.

An unsupervised ADMIXTURE analysis^26,27^ of the entire Western Eurasian dataset revealed different ancestral proportions among individuals from various Italian areas (Fig. 2C and S4). At K=4, which has the lowest cross-validation error, individuals from the LED and REN Valleys generally align with the latitudinal Italian pattern for three ancestral components, especially in LED_4GP and REN_4GP. An increased proportion of the silver component, which is linked to Neolithic expansions and maximized in the Sardinian population, is accompanied by decreased contributions of blue, which is likely associated with CHG/Iran_N ancestry, and green, which is probably of Middle Eastern origin and was introduced to Sardinia through Phoenician and Punic contacts. However, this latitudinal trend is not confirmed by the fourth component, which is maximized in populations from Russia (gold) and is likely related to EHG.

At higher Ks, 8 and 9 (Fig. 2C and S4), LED_4GP and REN_4GP individuals share and maximize a common ancestral component (red), while XGP retains a lower percentage. The same component is present in an individual from TAA (REN082), with grandparents from a locality between the Ledro and Rendena Valleys, whose genetic data was also placed between the two valleys in the Italian PCA (Fig. 2B). The maximization of a unique ancestral component is usually consistent with a history of relative genetic isolation and genetic drift, similar to patterns observed in populations such as Sardinians, BedouinB and Basques (at Ks 3, 4 and 6, respectively; Fig. S4).

From K=10 onward, an additional black component clearly differentiates REN individuals from LED ones (K10 in Fig. 2C and Fig. S4). This divergence across increasing K values suggests a common ancestral background followed by independent genetic drift in the two valley populations.

### Genetic signals of isolation and recent demographic pattern

In order to better understand the genomic patterns of the two valleys, which seems to point to a broader ancestral Alpine population with some micro-geographic peculiarities, we examined the distribution of runs of homozygosity (ROH)^28^, which quantify contiguous homozygous genomic segments inherited identically from both parents. Two complementary measures were considered: the number of ROH segments (*N*ROH) and the cumulative length of these segments (*S*ROH) across all Western Eurasian populations included in the dataset, with a focus on Italy. In general, smaller effective population sizes (Ne) result in more homozygous segments being inherited (*N*ROH), whereas mating among close relatives tends to increase their length and therefore the total *S*ROH. Notably, reduced Ne also raises the probability that two randomly chosen individuals share recent common ancestry.

Previous studies^7,11,12,14^ have shown that Sicilians and peninsular Italians behave as a panmictic population, whereas Sardinians represent a genetic isolate. Our ROH data (Fig. 3A) show that LED and REN have, on average, distributions similar to that observed in Sardinia, i.e. longer and more abundant ROHs. By contrast, North, Centre and South Italy, together with REN_XGP and LED_XGP individuals, show both a lower number of and shorter ROHs, consistent with a larger effective population size (Ne), greater admixture, or more exogamous mating patterns. We also examined the *S*ROH distributions across length classes for each individual: ROH segments were binned from 1.5 cM (minimum threshold chosen for reliable detection) to 4 cM and thereafter in 4cM intervals, with the final bin including segments >20 cM (Fig. 3B and S5A). Once again, our LED and REN individuals show a pattern similar to Sardinians, with most ROH segments longer than 4 cM (88.2% in LED_4GP; 96.2% in REN_4GP) (Fig. 3B), even higher than those observed in Sardinia (85.2%), and in stark contrast to other Italian populations (42.9%, 35.9% and 59.1%, for North, Centre and South Italy, respectively). Moreover, Sardinia, LED_4GP, and REN_4GP have significantly more individuals with high NROH and SROH with respect to North, Centre, and South Italy (Fig. S5B and S5C).

**Fig. 3.**
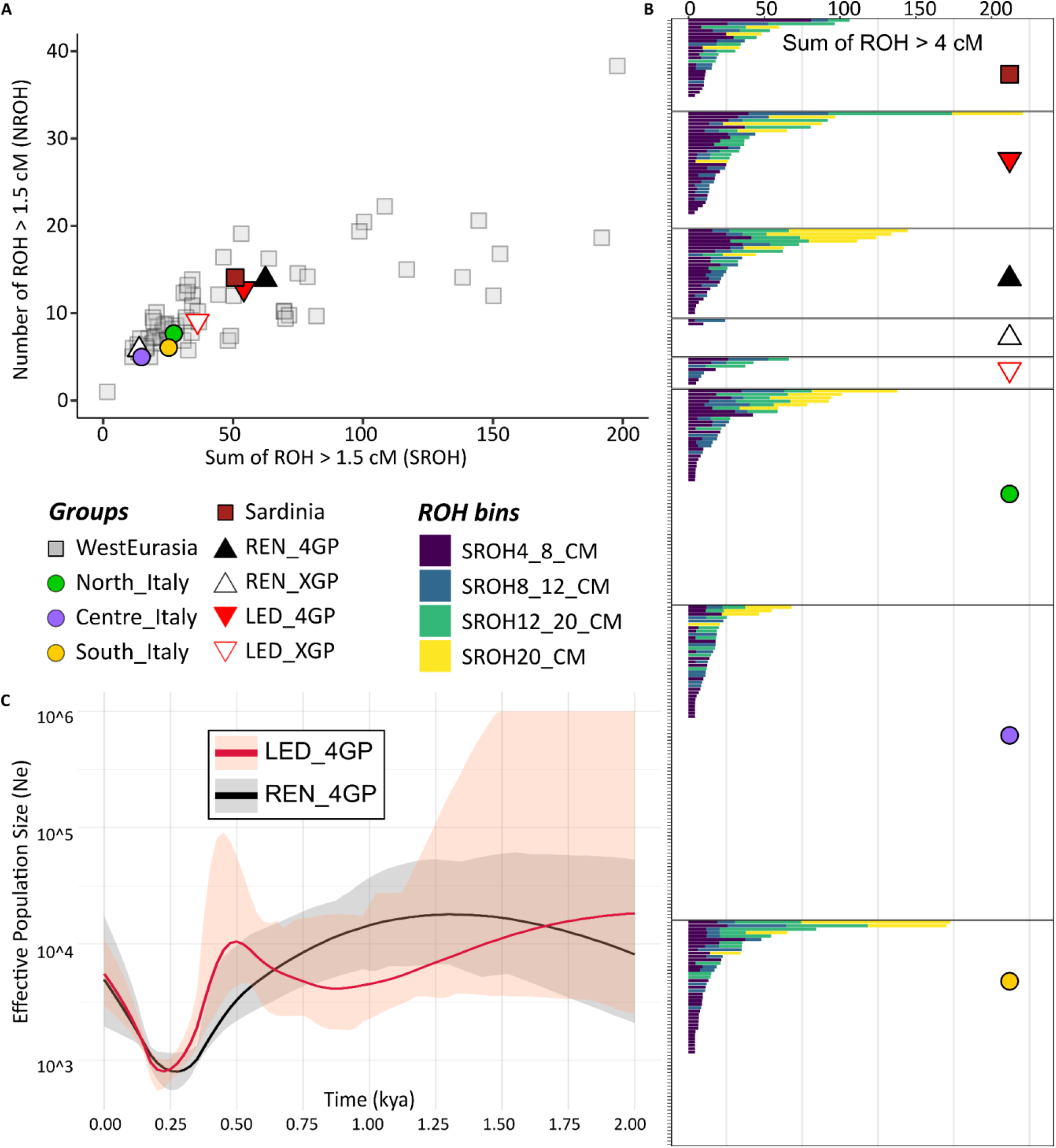
Tracing genetic signals of isolation and recent demographic histories. Legend at center-left of the figure. **A)** Mean SROH and NROH for all populations in Western Eurasia, ROH segments > 1.5 cM. **B)** SROH on ROH fragments > 4 cM, binned by length, in Italian populations. Individuals are shown on the y-axis, including those without ROH fragments over the threshold. **C)** Effective population size variation inferred from IBD segments for Ledro Valley and Rendena Valley in the last 2 kya. Solid line is the Ne estimation while shaded area represents 95% confidence interval. See also Fig. 1C for Ne comparison with mtDNA.

Given the elevated autozygosity observed in ROH analyses, we used within-population IBD segments to estimate changes in population size over the past 2 kya (Fig. 3C). In this time frame, a slight decrease in Ne is evident in both valleys, first in LED_4GP and then in REN_4GP about a millennium later, supporting the demographic trend inferred from the mitogenome-based BSP (Fig. 1C). Intriguingly, LED showed signs of recovery between one thousand and five hundred years ago, which could explain the plateau observed in the mtDNA BSP. Both valleys experienced a significant more recent bottleneck around 200-300 ya, followed by partial recovery. This timing coincides with the last historically documented plague epidemics in northeastern Italy, which also affected cattle in the Rendena Valley^29–31^. This recent bottleneck is not visible in the mtDNA BSP (Fig. 1C) since mitogenome datasets are more effective in older timeframes. On the other hand, the genome-wide data requires larger sample sizes and whole-genome sequencing data to inquire over longer periods.

### Genomic structure within the two Alpine valleys

We extended our haplotype analyses by using the ChromoPainter/fineSTRUCTURE^32^ tool. PCA based on haplotype sharing confirmed the distinct clustering of LED and REN individuals with respect to the Italian population (Fig. S6A). Among the first ten principal components, PC4 and PC5 (explaining 2.84% and 1.98% of the variance, respectively) separated these two Alpine valleys from the broader Western Eurasian variation, whereas PC10 (0.74%) distinguished LED_4GP and REN_4GP (Fig. S6B).

Due to the unique characteristics of the two valleys within the Italian genomic landscape, we used FineSTRUCTURE to search for potential population structures. LED and REN belong to different clusters of the same “modern Alpine” branch and are clearly separated from other Italian populations (Fig. 4A and S7). Additionally, they exhibit an unexpected internal structure: two distinct clusters among REN_4GP (named Rendena Valley A and B) and three for LED_4GP (Ledro Valley A, B and C).

**Fig. 4.**
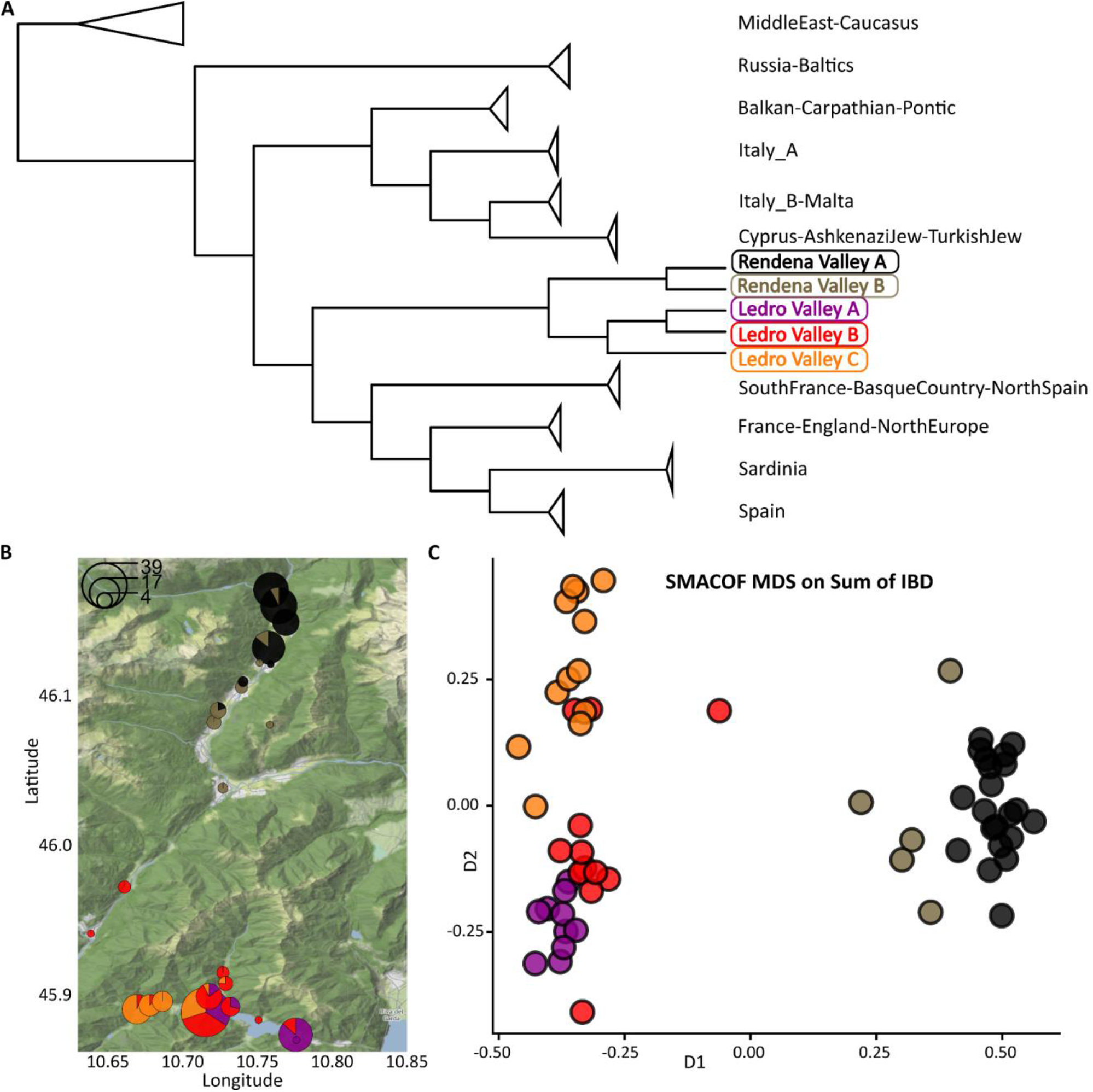
Geographic structure in the two Alpine valleys. **A)** ChromoPainter/fineSTRUCTURE cladogram of Western Eurasian populations, inferred excluding admixed individuals (XGP). Population clusters are grouped by geographic region to aid visualization. Branch lengths preceding triangles reflect the height at which the dendrogram was truncated; triangle sizes are proportional to the number of collapsed branches. See Fig. S7 for complete tree, Dataset S1 and S4 for cluster membership. **B)** Geographic distribution of the grandparents (GPs) of the analyzed individuals. Pie charts indicate cluster membership, with radii proportional to the number of grandparents originating from the same location. GPs reported generically as “Ledro” were placed at the confluence of the three Ledro sub-valleys (largest pie). Colours for alpine clusters as in A. See Dataset S1 for individuals’ coordinates. **C)** Multidimensional scaling (MDS; SMACOF algorithm) based on the sum of IBD sharing among individuals from LED and REN clusters. Same colours as in A and B. See also Fig. S8.

This genomic structure is strongly correlated with the microgeographical GP origin of each individual (Fig. 4B, Dataset S1). The two REN clusters, RendenaValleyA and RendenaValleyB, have statistically different latitudinal distributions (Bonferroni-corrected pairwise t-test, t = 6.68, p = 6.59×10⁻⁷). One cluster is mainly found in the north, and the other is mainly found in the south, reflecting the valley’s north-south geographical orientation. The overall spatial differentiation is also confirmed by PERMANOVA (R² = 0.644, F = 43.40, p = 0.0002). Instead, LedroValleyA, LedroValleyB, and LedroValleyC are primarily correlated with longitude (Bonferroni corrected pairwise t-tests: t = 5.43 and 3.20, p = 6.85×10⁻⁴ and 0.006), reflecting the east-west geographical orientation of the valley and the division in three subvalleys (Ledro, Ampola and Concei), again confirmed by PERMANOVA (R² = 0.695, F = 23.96, p = 0.0001). In both cases, beta-dispersion tests indicated no significant difference in within-group variability (REN: p = 0.50; LED: p = 0.49), supporting the validity of the between-group spatial differentiation.

The substructure found within the two valleys is also supported by SMACOF multidimensional scaling (SMACOF MDS) derived from pairwise identity-by-descent (IBD) sharing estimates between individuals (Fig. 4C and S8). RendenaValleyA and RendenaValleyB are separated along D1, while LED clusters are mainly distinguished by D2. Even though separation is not complete (especially for LedroValleyB), regardless of which specific sequences are shared, this analysis confirms distinct patterns of haplotype sharing between clusters, consistent with the ChromoPainter/fineSTRUCTURE results.

### Genetic affinities with other Western Eurasian populations

We used f-statistics to test if the different genetic clusters identified within the two Alpine valleys show preferential affinities (i.e., asymmetric gene flow and shared genetic drift), relative to other Western Eurasians, with respect to the three major ancestral sources that contributed to present-day European populations^4,33^: Western Hunter-Gatherers (WHG), Anatolian Neolithic farmers (Anatolia_N) and pastoralists from the Pontic-Caspian Steppe (Yamnaya), as defined in the Dataset S6. To address this, we computed *f*4(Mbuti, modern Western Eurasian clusters; Anatolia_N, and either Yamnaya or WHG) (Dataset S7). All of the modern LED and REN clusters defined here exhibit consistent patterns that overlap with each other and are close to, yet distinct from, the other Italian clusters (Fig. 5A). This picture was clarified by including individuals from TAA from the Mesolithic to the Middle Bronze Age^19^ and other ancient Western Eurasian reference data^6,21,34–44^. Ancient TAA populations, modeled by qpADM^45^ as a mixture of WHG and Anatolia_N (including Ötzi)^19,20^, behave similarly to the Sardinia clusters. However, those ancient TAA groups populations that have been shown to include a Steppe component, CA_EBA_2 (Chalcolithic_EarlyBronzeAge_2) and MBA (MiddleBronzeAge) by Croze et al. 2025^19^, are much closer to our modern Alpine clusters (Fig. S9). The genetic affinity of modern LED and REN clusters with Yamnaya-related populations is further supported by the high frequency (74%) of Y-chromosome haplogroup R1b, detected in 26 individuals with their terminal paternal ancestor (TPA) from one of the two valleys, a lineage commonly associated with the demographic expansion of Yamnaya-related groups during the Bronze Age (see Dataset S1 and Supplementary Text)^46^ and that is more frequent in northern than southern Italy^47^.

**Fig. 5.**
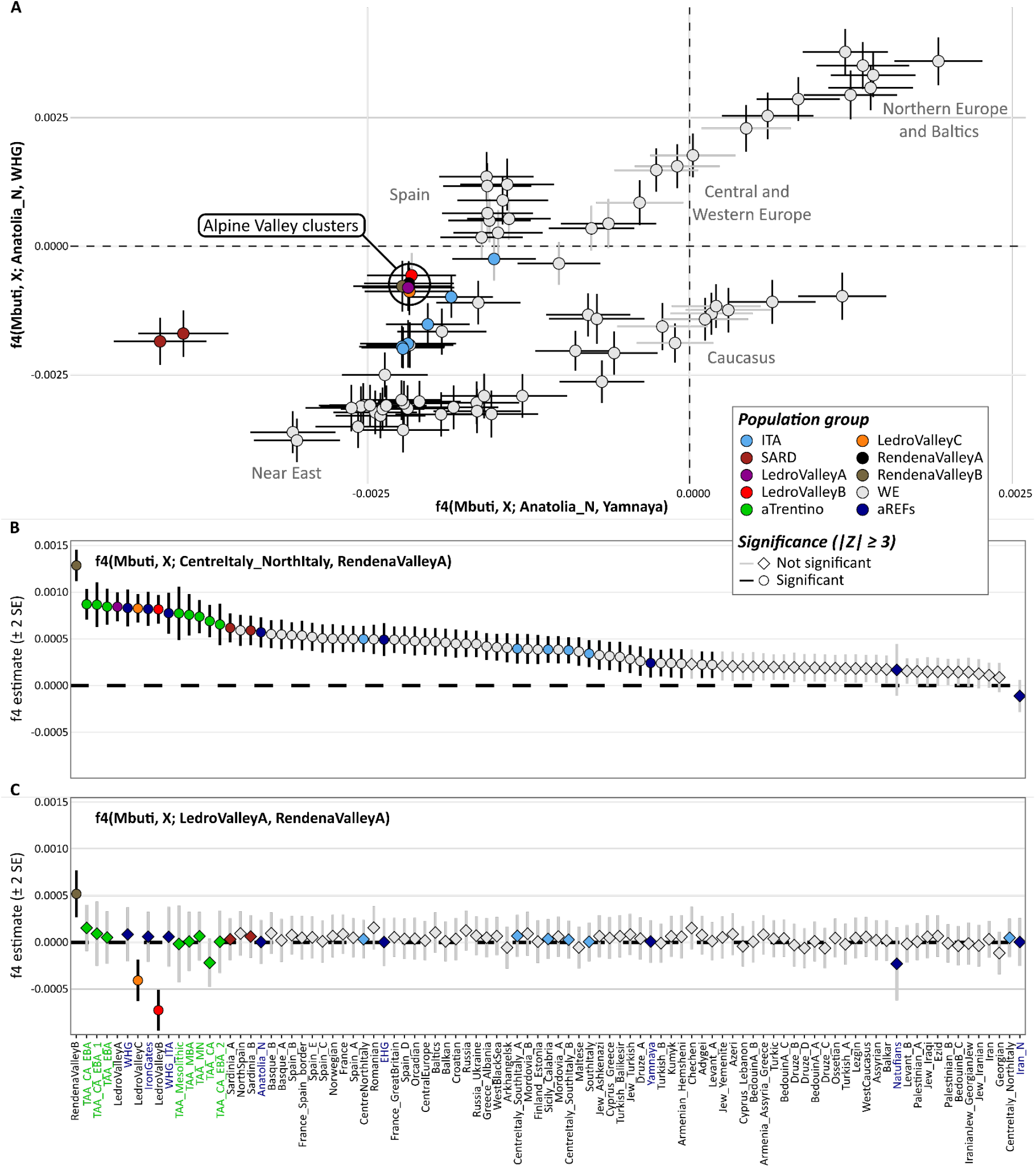
Genetic connectivity in eastern Alpine valleys and affinities with Western Eurasia. **A)** Affinities of Western Eurasian clusters toward Anatolia_N relative to either Yamnaya or WHG. Alpine clusters show closest affinity to other Italian populations. Error bars represent ±2 standard errors. Legend in the inset. **B)** f4 statistics testing the treeness of Western Eurasian clusters plus selected ancient populations relative to a representative Alpine valley cluster (RendenaValleyA) and the closest Italian cluster (CentreItaly_NorthItaly). Legend as in A, except for “aTrentino” that refers to ancient TAA plus the following suffixes: CA: Chalcolithic; EBA and MBA: Early and Middle Bronze Age. “aREFs” that are ancient populations thought to have influenced Western Eurasia genetic makeup. **C)** Testing differential affinities, with the same populations as in B, between Ledro Valley versus Rendena Valley. Legend as in A and B, same order of populations as in B.

Since modern Alpine and CentreItaly_NorthItaly clusters have similar results for *f*4(Mbuti, X; Anatolia_N, and either Yamnaya or WHG), we tested their differential affinities with other modern and ancient populations, using RendenaValleyA as a proxy (Fig. 5B). In *f*4(Mbuti, X; CentreItaly_NorthItaly, RendenaValleyA), a clear hierarchical pattern emerges. The strongest affinity is between the two REN clusters, followed by a set of population groups with comparable affinities, including the LED clusters, ancient TAA individuals and WHGs, and then Sardinians, Basques and Anatolia_N. All other Western Eurasian modern populations and ancient reference groups display lower affinities. Finally, we tested whether Western Eurasian populations display preferential affinity toward one valley over the other. No population groups showed significant differences, except for the clusters from the same valley (Fig. 5C). This suggests that the differences in genomic makeup between LED and REN clusters are not due to differential admixture or shared drift with external groups.

Outgroup f3-statistics mirror the patterns observed with f4-statistics (Fig. S10, Dataset S7). LED and REN clusters overlap closely with each other and with Central/Northern Italian groups, while also showing slightly increased shared drift with Anatolia_N relative to other Italian and Western European populations (Fig. S9A and S9B). In both comparisons, they occupy an intermediate position between Sardinians/ancient TAA groups and Western Europeans. In contrast, Basque and Spanish populations show relatively higher shared drift with WHG. Directly comparing LedroValleyA and RendenaValleyA reveals largely symmetric patterns of shared drift across all populations except for the valley-specific Alpine clusters (Fig. S10C).

## Conclusions

The eastern Italian Alps have historically been a key geographic corridor for cultural and genetic exchange between Mediterranean and Central European populations. Paleogenomic inferences have mostly relied on the high-coverage genome of the Copper Age Tyrolean Iceman who lived around 5300 ya and exhibited an exceptionally high proportion of Anatolian Neolithic ancestry with low but detectable WHG-related ancestry. A recent archaeogenomic study proved that this genetic profile is typical of the region and the Yamnaya-related ancestry emerged in the region around 4400 ya^19,48^. Surprisingly, this level of genomic detail has never been reached in modern populations of the Alpine valleys to understand if and to what extent they have retained their ancient genomic makeup.

To assess this issue, we analyzed the genomic variation of the contemporary communities in the Ledro and Rendena Valleys using complete mitochondrial genomes and genome-wide SNP data within a novel Italian dataset. Our findings show that the modern LED and REN populations form a distinct “modern Alpine” genomic group, clearly different from the broader Italian pattern of variation. Individuals with four grandparents from these valleys share a genetic background characterized by higher levels of Neolithic farmer ancestry than non-Sardinian Italians, similar to modern Sardinians and in continuity with ancient local populations, including the Tyrolean Iceman^19,21^.

The long-term evolutionary history of the two valleys is well summarized by the mtDNA data, which is more informative over long time frames. Variation in effective population size based on mitogenomes showed a sharp increase during the early Neolithic period. Over the last two millennia, isolation and subsequent genetic drift have differentially affected the two valleys. This pattern is explained by the redundancy of many haplotypes, which points to isolation events splitting the shared ancestral metapopulation into two genetically distinct valley communities, as further testified by distinct, valley-specific mtDNA haplogroups and newly defined sub-haplogroups.

At the genome-wide level, the history of relative geographic isolation and independent genetic drift is attested by higher levels of autozygosity (runs of homozygosity), similar to those of other isolated populations, such as Sardinians, in contrast to the rest of the Italian peninsular modern populations.

Estimates of population size variation using IBD segments, which are more effective within a two-millennium timeframe when using SNP panels^49^, confirm asynchronous decreases in both valleys over the past two thousand years, first in Ledro Valley and then in Rendena Valley about a millennium later. More recently, IBD data highlights a sharp, synchronized population bottleneck approximately 200 to 300 years ago, coinciding with documented plague outbreaks that potentially contributed to demographic decline.

Remarkably, this genetic isolation and structured drift persist at an extremely fine microgeographic scale within the valleys themselves. Despite their similarities, a high-resolution analysis of haplotype sharing reveals an internal substructure that directly correlates with local topography and specific grandparental geographic origins. The Rendena Valley exhibits distinct fine-scale genetic differentiation along a north-south axis, reflecting the valley’s orientation. In contrast, the Ledro Valley resolves into three genetic subclusters aligned with its three subvalleys (Ledro, Ampola, and Concei) along an east-west axis. We hypothesise that this differentiation has been caused by geographic barriers, limited mobility and local marriage practices favoring endogamy, potentially linked to the preservation of land ownership, livestock resources or community structure. In the Rendena Valley, the same practices might have contributed to local linguistic heterogeneity. Similar tendencies could be present in other TAA Valleys; unfortunately, no comparable genomic data from other present-day alpine individuals are currently available, which prevents any possible comparison or validation of this hypothesis.

These modern LED and REN clusters exhibit overlapping patterns and a significant excess of shared drift related to Anatolia Neolithic compared to WHG and Yamnaya. Their behavior is similar to, yet distinct, from that of other modern Italian groups. The valleys’ genetic heritage is deeply tied to the region’s history, at least since the Bronze Age. In fact, they share a much similar pattern with Early and Middle Bronze Age clusters of ancient individuals with Yamnaya-related ancestry than with older regional individuals lacking such an ancestry, such as the Tyrolean Iceman^19,20^. External Western Eurasian populations show no preferential affinity for one valley over the other, and significant differential affinities are restricted entirely to clusters within the same valley. This suggests that, despite their shared regional roots, the modern inhabitants of the Ledro and Rendena Valleys possess a distinct and independent genomic makeup, inherited by peculiar ancestral gene flows from outside and later shaped by localized genetic drift.

In conclusion, for the first time, our findings bring the genomic history of two eastern Alpine valleys up to the present day. We demonstrated how geographic barriers and localized endogamy can preserve ancient genomic components and create highly localized, fine-scale genetic structures over centuries, even among neighboring Alpine communities.

Future work should expand these analyses to other Alpine valleys, and include high-coverage whole-genome sequences. This will enable finer demographic inference, as well as the study of signals of natural selection and/or medically relevant variants. Most importantly, to reconstruct the deeper population history of the eastern Italian Alps, it will be crucial to integrate ancient DNA from local archaeological contexts, such as the Ledro pile-dwelling site.

## Materials and methods

### Sample collection

#### Sample acquisition

We collected 121 saliva samples through mouthwash rinsing from participants living in Rendena Valley, and 141 saliva samples from Ledro Valley inhabitants, both areas are located in TAA. REN individuals were sampled in the municipalities of Carisolo, Caderzone and Vigo Rendena, while the LED sample collection took place in Molina di Ledro, in accordance with relevant guidelines and regulations. Written informed consent was obtained from all donors, who also provided information about their place of birth and geographical origins, as well as genealogies traced back up to their great-grandparents. Following the same procedures, we also collected 139 saliva samples from Italians with diverse regional origins.

#### Ethical approval

All DNA samples and genealogical data analyzed here were collected on a voluntary basis after obtaining appropriate informed consent. All experimental procedures were approved by the Ethics Committee of the University of Perugia (protocol numbers 2017-01), revised on October 18^th^, 2023. The use of the “historical” collections in genomic studies was also approved by the Ethical Committee Fondazione IRCCS Policlinico San Matteo in Pavia (protocol number 0028298/22).

### From extraction to screening and mitogenome sequencing

#### DNA extraction

Automated DNA extraction from saliva samples was performed using the Maxwell RSC Blood DNA kit, deemed suitable also for saliva samples.

#### MtDNA-guided screening

Complete mitogenome sequences were obtained through Next Generation Sequencing (NGS) using the Illumina MiSeq platform. Library preparation was carried out following the Illumina DNA Prep protocol v10. After quality assessment with FastQC^50^, the resulting FASTQ files were trimmed using Trim Galore v0.6.4_dev^51^ to remove adapter sequences and low-quality bases (Phred score <30) at read ends. Reads were aligned to the revised Cambridge

Reference Sequence (rCRS; NC_012920.1)^52^ using the BWA mem algorithm^53^. The resulting BAM files were filtered and sorted with SAMtools^54^, and duplicate reads were removed using Picard’s *MarkDuplicates* tool^55^.

Downsampling and variant calling were performed using the mtDNA-Server v2^56^ in fusion mode, which integrates Mutserve2^57^ and Mutect2. Variants were detected using a minimum variant allele frequency (VAF) threshold of 0.1. Reads were downsampled to a maximum coverage of 2000×, retaining only positions with a minimum mean coverage of 10×. Additional filtering parameters included a minimum base quality of 30 and a minimum mapping quality of 30.

Final haplotype and haplogroup assignments were performed using Haplogrep v3.2.1^58^. When diagnostic haplogroup-defining positions were flagged as missing in the Haplogrep report, we manually inspected the corresponding BAM files to verify their presence. Individuals with successful DNA extraction and sequencing were further filtered using stringent quality criteria: absence of contamination, average depth ≥10×, genome coverage ≥90%, less than three heteroplasmies per individual and terminal maternal ancestor provenance. After applying these filtering criteria, 94 mitogenomes from Val Rendena were retained out of 116 sequenced individuals, while 91 mitogenomes from Valle di Ledro were retained out of 100 sequenced individuals. Sequenced individuals with a TMA originating from regions other than the two valleys were included in the mtDNA comparative analyses with Western Eurasian populations.

#### MtDNA analyses

Haplotype diversity (Hd) was calculated as a measure of the probability that two randomly chosen haplotypes from the population are different, based on the observed haplotype frequencies. Nucleotide diversity (π), defined as the average number of nucleotide differences per site between all possible pairs of sequences in the sample, was computed using the ape package v4.3.2^59^ in R v4.3.2^60^. The calculation was performed on aligned mitogenome sequences, considering pairwise differences across all sites. We performed the multiple sequence alignment with software MAFFT with option *--auto*^61^.

Individuals were grouped according to their macro-haplogroup and geographical origin. Haplogroups observed in at least four individuals were defined as macro-haplogroups. Haplogroups with lower frequencies were instead aggregated into broader categories according to their cladistic relationships. When extending the comparison of Rendena and Ledro Valleys to the broader contexts of northeastern Italy and Western Eurasia, and applying the same classification criteria, additional macro-haplogroups were defined.

PCA was performed using the package FactoMineR^62^ in R^60^. A chi-square test on the contingency table of the distribution of macro-haplogroups was performed also using R. After removing all positions containing gaps and ambiguous data, a MP tree was built with mtPhyl v.5.003^63^ and then manually corrected using Haplogrep 2.4^64^ phylogenetic trees. Coalescence ages of haplogroups and subhaplogroups were calculated on the trees by means of the average number of base substitutions (rho-ρ) in the mtDNA, disregarding “indels,” hot spots, the transition at nucleotide position (np) 16519 and variants around np 309 (from np 303 to np 315) and np 16189 (from np 16182 to np 16194). Standard error (sigma-σ) was calculated from an estimate of the genealogy^65^. Subsequently, we converted the rho and sigma parameters into years as indicated by Soares et al. 2009^66^.

Further time estimates and demographic trends were evaluated using Bayesian Evolutionary Analysis of Sampling Trees (v2.7.7)^67^. The Hasegawa-Kishino-Yano substitution model with gamma-distributed rate variation among sites (HKY + G) was selected as a common model for both datasets, as it has been widely adopted in human mtDNA studies, including Sardinians and other isolated populations^13,68^. The transition/transversion ratio was estimated during the run, and the gamma distribution was discretized into eight categories to account for rate heterogeneity. A strict molecular clock was applied, assuming a constant substitution rate across all lineages. A coalescent Bayesian Skyline model was chosen as a prior to allow for fluctuations in past population size. The clock rate followed an informative normal prior with a mean of 2.45 × 10^−5^ and a SD of 3.5 × 10^-6^ ^66^. Haplogroup assignments for each sequence and the corresponding haplogroup age estimates, based on previously published divergence times^69^, were incorporated as Bayesian priors and modeled using a normal distribution. The Markov chain Monte Carlo chain was run for 10 million generations. Tracer v1.7.2^70^ was used to assess effective sample sizes and generate the BSP to visualize past demographic trends, together with the package ggplot2^71^ in R^60^.

### Genotyping and dataset curation

#### Genotyping

Axiom^TM^ Genome-Wide Human Origins 1 Array^72^ (HO SNP chip) (629,443 SNPs), was used to genotype 48 individuals sampled in Ledro Valley and 48 in Rendena Valley. After filtering for missingness (<2%) and call rate (>95%), we excluded two individuals from LED. From individuals sampled in Rendena Valley, we added three individuals to the Italian dataset: one from TAA and two from the Lombardy region. Two individuals were classified as ADMIXED and two with Trentino_3GP, with only three grandparents (GP) from the region (See Dataset S1A).

#### LED dataset

We divided the final 46 LED individuals into groups according to their genealogies. This resulted in 36 LED individuals with all four GPs from the Ledro Valley (LED_4GP), six with three GPs (LED_3GP), one with two GPs (LED_2GP) and four with 1 GP (LED_1GP).

#### REN dataset

We also divided the 41 genotyped REN individuals into groups based on their genealogies, obtaining 29 REN individuals with all four GPs from Rendena Valley (REN_4GP), seven with three GPs (REN_3GP), four with two GPs (REN_2GP) and one with one GP (REN_1GP).

#### Our Italian dataset

Following the same procedure, we genotyped 139 Italian individuals using the HO SNP chip, in addition to three Italians sampled in Rendena valley (See Dataset S4). After filtering for missingness (<2%) and call rate (>95%), we excluded two individuals from Latium, Italy. These individuals originate from all regions of Italy and, together with publicly available data, ensure a minimum of four individuals per region (label: RegionThisStudy). All individuals have four GPs from their assigned region, either from the same province within that region, or with at most one GP originating from a different province.

#### Modern Western Eurasia dataset (Modern WE)

We selected Human Origin genotyped modern Western Eurasian populations (WE; defined as countries not on the African continent with longitudes between −15 and 60 and latitudes higher than 22; populations chosen as in Aneli et al. 2021^5^, plus Western Eurasian jewish diaspora populations) from AADRv62^22,23^ (See Dataset S5).

#### Modern Mbuti

In order to perform f-statistics with a divergent clade, we also selected Mbuti.HO from Sub-Saharan Africa, as in^19,20,73^ (See Dataset S5).

#### Ancient Western Eurasia dataset (Ancient WE)

Following the same procedure for the Modern WE dataset, we selected from AADRv62^22,23^ a subset of ancient individuals (haploid genome (AG) captured on 1240k SNP chip) with the “ASSESSMENT” column equal to “PASS”, and not marked as outliers (“_o”) or as related (“_1d” and others), choosing populations that are likely to be source of current variability inside Western Eurasia based on previous publications^6,8,73–75:^ Western Hunter Gatherer (WHG), WHG_Italy (WHG_ITA), “IronGates”, Iran Neolithic (Iran_N), Anatolia Neolithic (Anatolia_N), “Yamnaya” and “Natufians”. In addition, we used all data from ancient TAA individuals in Croze et al. 2025 and Müller et al. 2003^19,20^ (TrentinoAltoAdige_ plus their final attribution in the study, e.g. Copper Age - Early Bronze Age in TAA is here TrentinoAltoAdige_CA_EBA, see Dataset S6). We also used shotgun sequencing data for the Tyrolean Iceman individual, due to his importance as a reference for Alpine population history.

#### Filtering and kinship for all modern individuals

After filtering for SNP missingness (<2%) inside each modern dataset, we removed three Italians from the Italian dataset. After kinship detection inside REN population, we removed four individuals (3 REN_4GP, 1 REN_3GP) due to relatedness from first to third degree (relatedness detected with PLINK2 v2.0.0-a.6.9LM^76^, option–make-king-table^77^, excluding samples with KINSHIP>0.0442) to obtain the final REN dataset. We applied the same approach to the LED dataset, removing one LED_4GP, one LED_3GP and one LED_1GP. The same procedure was applied across all Italian regions, and then to the combined Italian dataset, removing an additional four individuals. After merging with modern WE Dataset (creating complete Modern WE dataset), we checked for kinship within and across populations, eliminating 21 individuals.

### Allele-frequency based analyses

#### PCA on Italy and WE datasets

We filtered the complete modern WE dataset using PLINK v1.90b6.21^78,79^ using options --geno 0.02, --mind 0.02, --maf 0.01. PCA was calculated using smartpca v18140 from the EIGENSOFT package^80,81^ with lsqproject:YES and numoutlieriter:0 options, either giving the population list of Italy or of the whole WE.

#### ADMIXTURE

To perform ancestry analysis, we applied the same filters used for PCA to the complete modern WE dataset, cutting for linkage disequilibrium (LD) by using the flag –indep-pairwise 100 25 0.2, as in Raveane et al. 2019^7^. ADMIXTURE v1.3^26^ was run from K1 to K16, with 10 replicates for each K using default options, except for -s time and -j8. The best K was chosen based on a cross-validation error pattern. Results were plotted with PONG^26^.

#### *f*-statistics

Once the modern WE individuals were subdivided according to ChromoPainter/fineStructure clusters (see subsequent section “Haplotype based analyses”), we first merged the complete Modern WE dataset with modern Mbuti (then filtered with PLINK v1.90b6.21^78,79^ --geno 0.02 and --mind 0.02), and then added the Ancient WE dataset (--geno 0.6 and --mind 0.95). All *f-*statistics were performed with the software ADMIXTOOLS 2 v2.0^82^ in R^60^ with default parameters (Dataset S7), then plotted using tidyverse^83^ and patchwork^84^ packages.

#### Haplotype based analyses

We extended and integrated the results of the SNP frequency-based analyses to the haplotype level, by using the ChromoPainter/fineSTRUCTURE^32^ tool and identity-by-descent (IBD) analyses. Haplotypes were inferred from Western Eurasians’ genotypes by phasing with SHAPEIT v5.1.1^85^. We then used ChromoPainter on our phased dataset (excluding REN_XGP and LED_XGP individuals) to characterise patterns of haplotype sharing among individuals. The outputs of ChromoPainter were then used for two downstream analyses, PCA and fineSTRUCTURE.

#### Phasing

We phased the complete modern WE dataset (previously filtered with PLINK v1.90b6.21^78,79^, options --geno 0.02 and --mind 0.02) using SHAPEIT v5.1.1^85^. The input VCF files were produced from PLINK binary files and converted using bcftools^54^. Phasing was performed per chromosome, using the corresponding genetic map (build 37) provided by SHAPEIT5. Phased BCF files were converted using bcftools^54^ into VCF.

#### ChromoPainter

Phased VCF dataset was restricted by excluding samples from LED and REN with less than four grandparents from their respective valley, then converted using bcftools^54^ convert -- hapsample to .hap format. The resulting .hap.gz files were then decompressed and processed using the impute2chromopainter2.pl script to produce files in ChromoPainter^32^ format. A recombination rate file was generated and then corrected using a custom R script (to perform local linear interpolation, as in Moreno-Mayar et al. 2024^86^ to ensure consistency between the SNP list and the map. To estimate the switch and mutation rates (Ne and Mu) required by ChromoPainter, we selected five random chromosomes (3, 7, 13, 19, 23) and performed an expectation-maximization (EM) run using ChromoPainterv2, parameters -in -iM -i 10. Parameters inferred were respectively 208.595 (Ne) and 0.000358 (Mu), which were then used to run the final ChromoPainter analysis.

The final chunkcounts matrix was then normalized and PCA was calculated using prcomp() in R^60^, then graphed using tidyverse package^83^.

#### FineSTRUCTURE

The chunkcounts matrix obtained from ChromoPainter was used as an input for fineSTRUCTURE^32^. We ran an initial MCMC with one million burn-in iterations (-x 1000000), followed by one million sampling iterations (-y 1000000) thinned every 10,000 steps (-z 10000). A second MCMC run was then performed with the same number of sampling iterations but without further burn-in (-x 0). This step allows refining the posterior distribution and improving stability. We inferred the clustering tree using the Tree mode of fineSTRUCTURE based on the second MCMC. The .tree.xml file was used to extract summary statistics including the mapstate matrix (population-level relationships), options -X -Y -e X2, and the mean coincidence matrix (individual-level co-clustering probabilities), options -X -Y -e meancoincidence. The algorithm identified 99 clusters. However, to improve interpretability and robustness, the final number of clusters was reduced to 79 by merging groups with fewer than five individuals and/or with Total Variation Distance (TVD) < 0.03, a threshold commonly used to denote similarity in copying vectors^87^. The final tree was visualized using iTOL v7.5.1^88^. Geographical coordinates were determined as the mean between the coordinates of all four grandparents’ locations and correlation (denomination “Ledro” was set at the geographical center of the valley), with ChromoPainter/fineSTRUCTURE LED and RENgroups, was estimated using PERMANOVA^89^ in R, package vegan^90^; final map with pies was obtained using packages scatterpie^91^ and ggmap^92^.

### ROH and IBD

#### ROH

We filtered the complete Modern WE dataset with PLINK v1.90b6.21^78,79^ options --geno 0.02 and --mind 0.02. We scanned genomes for ROH by using PLINK (same parameters as Ceballos et al. 2018^28^) with --homozyg-snp 50, --homozyg-kb 300, --homozyg-density 50, --homozyg-gap 1000, --homozyg-window-snp 50, --homozyg-window-het 1, --homozyg-window-missing 5, --homozyg-window-threshold 0.05. We considered for our analyses fragments over 1.5 cM and we binned SROH in 1.5 to 4, >4 to 8, >8 to 12, >12 to 20 and >20, the last four categories as also done in Ringbauer et al. 2021^93^. Statistical analyses were performed in R v4.3.2^60^. Kruskal–Wallis and Dunn’s post hoc tests were conducted using the rstatix^94^ package, and figures were generated using ggplot2^71^ and ggpubr^95^.

#### IBD: MDS and SMACOF

To further verify the substructures found within LED and REN (see “Haplotype based analyses”), we identified IBD segments in Western Eurasia with hapIBD^96^, then created distance matrices, based on the count, the sum and the average of IBD segments shared. We used MDS and SMACOF^97^ multidimensional scaling, a MDS that preserves the order of distances instead of absolute numbers, thus avoiding skewing by outliers. We called IBD on the phased Modern WE dataset (see “Haplotype based analyses”, section Phasing) with hapIBD v1.0 15Jun23.92f^96^ using PLINK maps (GRCh37) provided with the software. From the output, we summarized, in R^60^, the IBD sharing between each pair of individuals in a pairwise matrix each, according to three metrics: total sum of shared IBD segment lengths (cM), total count of shared segments, and average segment length (cM). Analyses were restricted to individuals from LED and REN clusters identified with ChromoPainter/fineSTRUCTURE (see section “Haplotype based analyses”). Each similarity matrix was transformed into a dissimilarity matrix using two approaches implemented in the R package smacof^97^: a reciprocal transformation (sim2diss, method = “reciprocal”), used for standard interval MDS, and a rank-based transformation (sim2diss, method = “rank”), used for non-metric ordinal MDS (SMACOF). Multidimensional scaling was then performed using the mds() function, with type = “interval” for standard MDS and type = “ordinal” for SMACOF, as in Fibla et al. 2023^98^.

#### IBD: demographic inference

IBD segments were detected for each chromosome arm, following the recommendation of Fournier et al. 2023^99^ to avoid centromeric regions. To this end, phased VCFs were split by chromosome arm using the HapNe utility split_vcf, with automatic minor allele frequency (MAF) filtering at 0.01 and genome build GRCh37. hapIBD was then run independently on each chromosome arm, using the corresponding PLINK GRCh37 genetic maps. IBD and HBD output files were concatenated per arm, and adjacent segments were merged as described above. Effective population size (Ne) through time was estimated with IBDNe version ibdne.23Apr20.ae9^49^, run using a minimum segment length of 4 cM, as recommended for SNP array data.

#### Y-chromosome haplogroup inference

To characterize paternal lineages, we analyzed the Y-chromosome SNP content of the Human Origins genotype data from all 42 male individuals who passed quality control. Of these, 14 individuals have a terminal paternal ancestor (TPA) from LED, 21 from REN, and the remaining seven individuals have TPAs originating outside these valleys (Dataset S1). Starting from the PLINK files of the full dataset, we extracted male individuals and Y-chromosomal positions, yielding a dataset of 2,070 Y-chromosomal SNPs with an overall genotyping rate of 99.9%. Y-chromosome haplogroup assignment was performed with Yleaf v4.0^100^, using the PLINK input mode, which allows direct inference from SNP-array genotype data. The reference genome was set to hg19, consistent with the genome build used throughout this study. Haplogroups were predicted against two independent reference trees: the YFull phylogeny v14 (https://www.yfull.com/tree/) and the ISOGG tree (https://www.isogg.org/tree). Both classification systems are reported. For each sample, Yleaf evaluates the set of informative Y-SNPs covered by the array and assigns the most likely haplogroup based on the derived allele pattern; the number of valid markers supporting the assignment is also provided. Haplogroup assignments are reported as returned by the software. Although the HO SNP chip contains sufficient Y-chromosomal markers for reliable haplogroup assignment, the limited number of downstream informative SNPs often results in paraphyletic classifications (e.g., R1b1a1b1a1a*).

## Acknowledgements

We thank Simone Floresta, Manuela Pernter, Sabrina Buscè, Marta Gobbi, Amedeo Luigi Zanetti, Valentina Miretti, Erika Vitali, and all the participants that donated their DNA and institutions that helped bring this project to fruition. We acknowledge local administrations, in particular the Comune di Carisolo, for their logistical support.

## Funding sources

This work was supported by the Italian Ministry for Universities and Research (MUR) PRIN 2022 grants 2022SNEBJY (to A.To., H.L. and H.C.H.) and 2022Y8BSAL (to A.A.); Fondazione Cariplo - Bando Giovani Ricercatori 2023, rif: 2023-1373 (to N.R.M.); Fondazione CARITRO, project entitled “Genesi” (to D.R., A.F., and A.A.).

## Authors contributions

Conceptualization: G.V., N.R.M., A.T., N.N., E.P., R.R., L.C., H.C.H., A.F., P.A.M., A.To., and A.A. Data curation: G.V., A.T., V.C., I.P. and G.B. Formal analysis: G.V., A.T., V.C., I.P. and G.S. Funding acquisition: N.R.M., N.N., E.P., D.R., H.C.H., H.L., A.F., A.To. and A.A. Investigation: G.V. and A.T. Methodology: G.V., N.R.M., I.C., P.A.M. and A.A. Project administration: A.S., A.To. and A.A. Resources: N.N., E.P., V.C., R.D.G., A.O., A.S., H.C.H., H.L., P.A.M., A.To. and A.A. Software: G.V., N.R.M. and A.T. Supervision: N.R.M., H.L., A.F., P.A.M., A.To. and A.A. Validation: G.V., N.R.M., A.T. and A.A. Visualization: G.V., N.R.M., A.T., G.S., V.N. and A.A. Writing-original draft: G.V., N.R.M., A.T. and A.A. Writing-review & editing: G.V., N.R.M., A.T., N.N., E.P., V.C., I.P., G.S., V.N., I.C., R.D.G., G.B., E.R., D.R., R.R., L.C., A.O., A.S., H.C.H., H.L., A.F., P.A.M., A.To., and A.A. G.V., N.R.M. and A.T. contributed equally to this work and share first authorship. A.A. and N.R.M. are the corresponding authors. All authors have read and approved the final version of the manuscript.

## Competing interests

The authors declare that they have no competing interests.

## Data availability

This study coanalyzes newly generated data with publicly available datasets. Public datasets used are GenBank^101^, IGSR^102^ and AADRv62^22,23^, with accession codes specified for each sample, together with source dataset and the original publication (Datasets S3-6).

Newly generated Human Origin genotypes will become available upon publication in Zenodo^103^ (Dataset S1 and S4). Mitochondrial fastQ data have been deposited at SRA^104^ as BioProject and are publicly available as of the date of publication (Dataset S1). Mitochondrial fasta data have been deposited in GenBank and will become available upon publication (Dataset S1). All data needed to evaluate the conclusions in the paper are present in the paper and/or the Supplementary Materials.

## Notes

### Competing Interest Statement

The authors have declared no competing interest.

