## SupplementaryText & Figures for "From prehistory to present-day: How isolation shaped the distinct genomic makeup of Italian Alpine valleys"

### Supplementary Information

**Text S1. Shared mtDNA lineages**

**Text S2. Valley-specific mtDNA lineages**

**Text S3. Y-chromosome lineages in Rendena and Ledro Valleys**

**Text S4. Additional info on genome-wide data**

**References**

**Supplementary Figures S1–S10**

#### Text S1. Shared mtDNA lineages

Phylogenetic analysis of haplogroup J1 highlights both shared and valley-specific lineages (Fig. 1B). Haplogroup J1 has an estimated age of 24,105 years ago (ya) (95% CI: 13,089–35,665 ya). Overall, ∼53% of the J1 sequences are from Rendena Valley, while 47% are from Ledro. A lineage exclusive to Ledro is represented by two sequences belonging to J1b1b1. In contrast, J1c3 represents a shared lineage, with an estimated age of 9,871 ya (95% CI: 3,009–17,013 ya) (Dataset S2). Five sequences belong to J1c3, all from Rendena Valley. Nine additional sequences define a newly proposed subclade, J1c3e2a, with an estimated age of 770 ya (95% CI: 0–1,885 ya), all sharing the T5201C transition. Among these, two sequences are from Ledro Valley and a cluster of seven sequences, four from Rendena Valley and three from Ledro Valley, share the same haplotype, supporting the phylogenetic definition of this subclade.

Another mitochondrial lineage shared between the two valleys is haplogroup T2, with an estimated age of 18,783 ya (95% CI: 9,111–28,920 ya). Overall, 26 sequences belong to this haplogroup, of which 65.4% are from Rendena Valley and 34.6% are from Ledro Valley. Most T2 sequences fall within subhaplogroup T2b, dated to 10,090 ya (95% CI: 4,094–16,296 ya), which is strongly widespread in Rendena Valley, where 16 sequences are observed, compared to 6 in Ledro Valley. Despite the high number of sequences, T2b is represented by only six distinct haplotypes with substantial redundancy in both valleys. Notably, one T2b haplotype is shared between the two valleys, suggesting inter-valley connectivity.

Haplogroup H5 further illustrates patterns of shared ancestry and local diversification, with an estimated age of 10,801 ya (95% CI: 4,469–17,365 ya). In this geographic context, two main subclades are observed, displaying contrasting valley-specific patterns. Subclade H5a includes five haplotypes, of which two are exclusive to Rendena Valley and one is shared between Rendena and Ledro valleys. In contrast, subclade H5t is exclusively detected in the Ledro Valley and includes a newly proposed subclade, H5t1, defined by the A8053G transition, highlighting a local diversification process within Ledro Valley.

An additional newly defined subhaplogroup is observed in both valleys within haplogroup HV1, which dates back to 21,399 ya (95% CI: 6,650–37,202 ya). A total of six sequences belong to this haplogroup and cluster into a novel subclade, here designated as HV1e. All HV1e sequences share a common set of seven transitions (C64T, A93G, T252C, A3221G, T15310C,

A15562G, C16292T), indicating their monophyletic origin and revealing another distinct lineage present in both valleys.

#### Text S2. Valley-specific mtDNA lineages

Haplogroup H3 is exclusively detected in the Rendena Valley population and dates back to approximately 10,939 ya (95% CI: 3,672–18,512 ya). A total of eight mitogenomes belong to haplogroup H3, of which four are assigned to a newly proposed subclade, here designated H3ap1, with an estimated age of 1,287 ya (95% CI: 0–3,071 ya) and defined by the A7364G and G15773A transitions, indicating recent local diversification within the Rendena Valley. The G15773A mutation results in a valine to methionine substitution at position 343 of cytochrome b (MT-CYB) and has a pathogenicity score of 0.201 according to APOGEE[1], falling within the "likely benign" range ($0.062 < \text{score} \leq 0.265$), although close to the upper threshold of this category.

Haplogroup J2 is detected exclusively in the Ledro Valley population and dates back to approximately 31,743 ya (95% CI: 19,513–44,556 ya). All six mitogenomes assigned to this haplogroup derive from the Ledro Valley, indicating a strongly localized distribution. Within J2, two sequences belong to distinct sublineages of J2a, whereas J2b1a is represented by three sequences sharing the same haplotype within Ledro, pointing to a local amplification of this subclade.

Finally, haplogroup U5a further emphasizes the asymmetric distribution of mitochondrial lineages between the two valleys. This haplogroup, with an estimated age of 18,014 ya (95% CI: 9,604–26,778 ya), is present almost exclusively in Rendena Valley and is represented by subclades U5a1 and U5a2. Subclade U5a1, dated to 16,172 ya (95% CI: 7,051–25,727 ya), includes eleven sequences from Rendena Valley and shows a pronounced redundancy, with haplotype U5a1a1 shared by seven individuals. Subclade U5a2 comprises six sequences, five from Rendena Valley and one from Ledro Valley. Notably, the sub-branch U5a2c1 is shared between the two valleys.

#### Text S3. Y-chromosome lineages in Rendena and Ledro Valleys

To complement the uniparental analyses, Y-chromosome haplogroups were inferred from male individuals in the HO dataset using Yleaf. Among the 35 individuals with a terminal paternal ancestor (TPA) within their respective valley (21 Rendena Valley, 14 Ledro Valley; Dataset S1), haplogroup R1b predominates in both populations (74.3%; Rendena: 71.4%, Ledro: 78.6%), consistent with high R1b frequencies reported in Northern Italy and with the West European Bell Beaker-associated Bronze Age dispersal[2]. Haplogroup J2 is the second most frequent, present in four individuals (11.4%; 3 Rendena, 1 Ledro), representing lineages of Near Eastern origin associated with Neolithic migrations into Europe[3]. Haplogroup E1b1b accounts for a further 8.6% (2 Rendena, 1 Ledro) and is widely distributed across the Mediterranean basin and North Africa, consistent with episodes of gene flow into the Italian Peninsula during historical periods[3]. A single individual from Rendena carries G2a2b, a lineage associated with the Neolithic expansion in Europe, and found in the Eastern Italian Alps from the Middle Neolithic through the Bronze Age[4]. One individual from Ledro carries haplogroup I. Haplogroup frequencies are broadly comparable between the two valleys, with no major differentiation in paternal lineage composition (full details in Dataset S1).

#### Text S4. Additional info on genome-wide data

In the western Eurasia PCA, the peculiar position of two individuals from Friuli Venezia Giulia region (North Italy), within Central European variability, could reflect the complex demographic history of this area, which includes several long-standing ethno-linguistic minorities. While all four GPs of these individuals were born within Italy, their genetic affinity with Central European populations may be consistent with ancestry linked to Slavic- or German-speaking communities historically present in the region.

Additionally, Sardinia variability is separate from the rest of Italian populations by PC2 (Fig. S3), while it is equidistant from both PC1 and PC3 (Fig. 2B). Considering the genealogical subgroups, LED_4GP and REN_4GP have internal axes of variability, even without considering LED_XGP and REN_XGP, which are drawn toward the rest of Italy according to their respective genealogies. Sole exception, two LED_XGP, with unknown genealogies except for TMA, seem to behave similarly to LED_4GP both in PCA and ADMIXTURE (Fig. 2). Moreover, one individual (REN082), from a nearby valley in Trentino (Giudicarie Inferiori), falls between the genetic variability of the two valleys, probably due to the geographical location.

#### Supplementary Figures

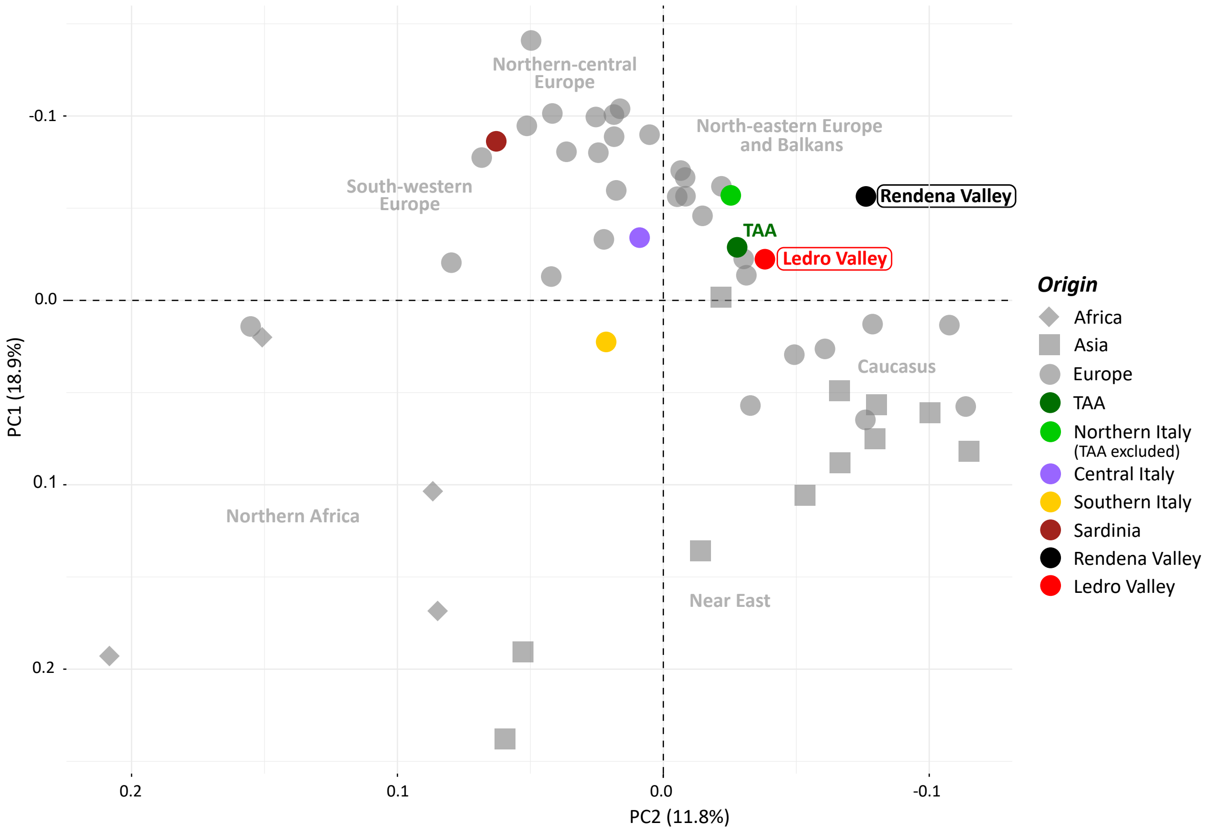


**Fig. S1. Principal Component Analysis (PCA) of mitochondrial lineage distributions in the Rendena and Ledro Valleys within the Western Eurasian context.** PCA based on mtDNA haplogroup frequencies (254 macro-haplogroups, Dataset S3) in the Western Eurasian context. “TAA” refers to Trentino-Alto Adige/Südtirol.

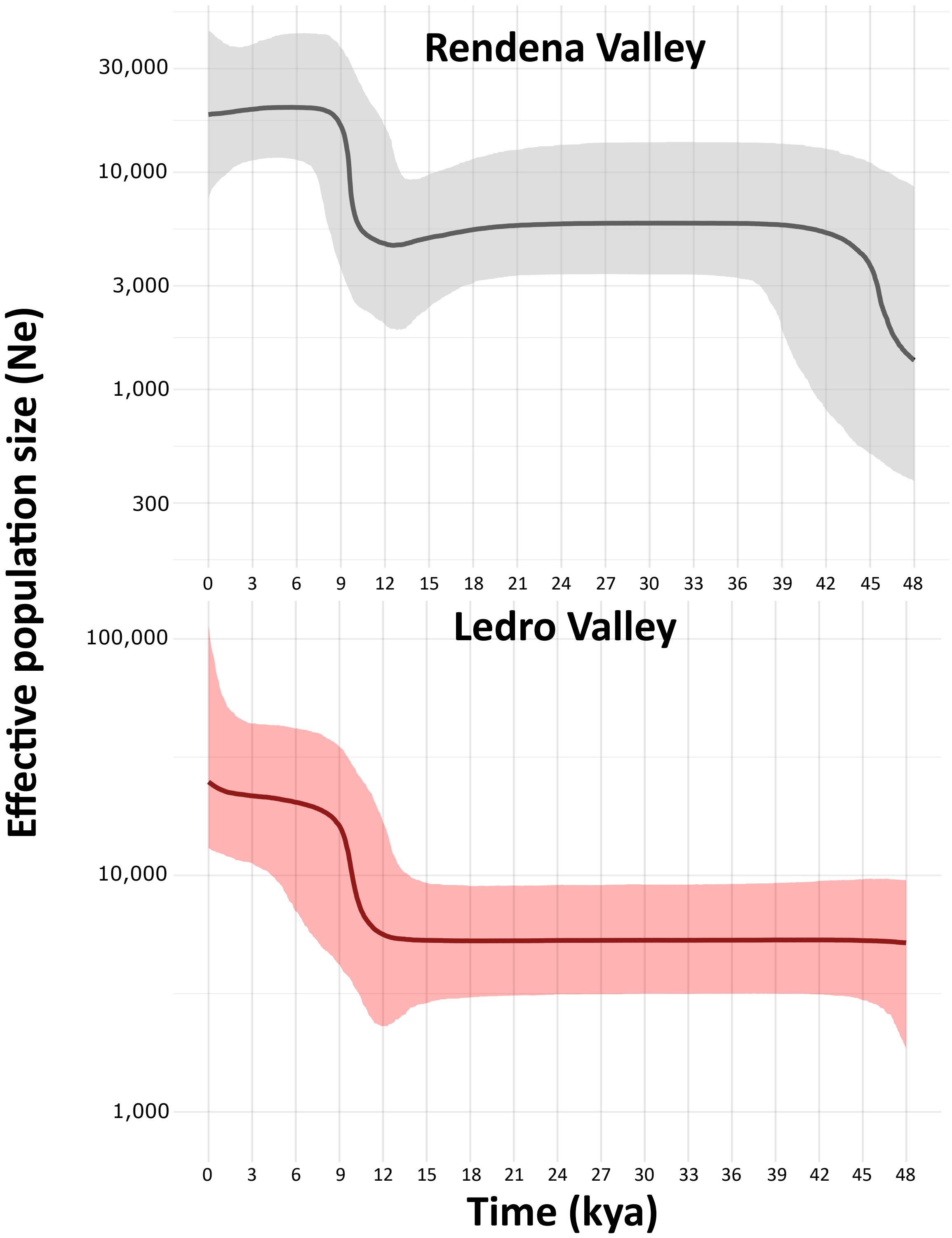


**Fig. S2. Bayesian Skyline Plots (BSPs) of Rendena and Ledro Valleys based on unique haplotypes.** BSPs were reconstructed by considering each haplotype only once. Compared to the full dataset, the reduced signal of decline in effective population size suggests that haplotype redundancy contributes substantially to the inferred demographic pattern. Analyses were performed using 1500 time intervals.

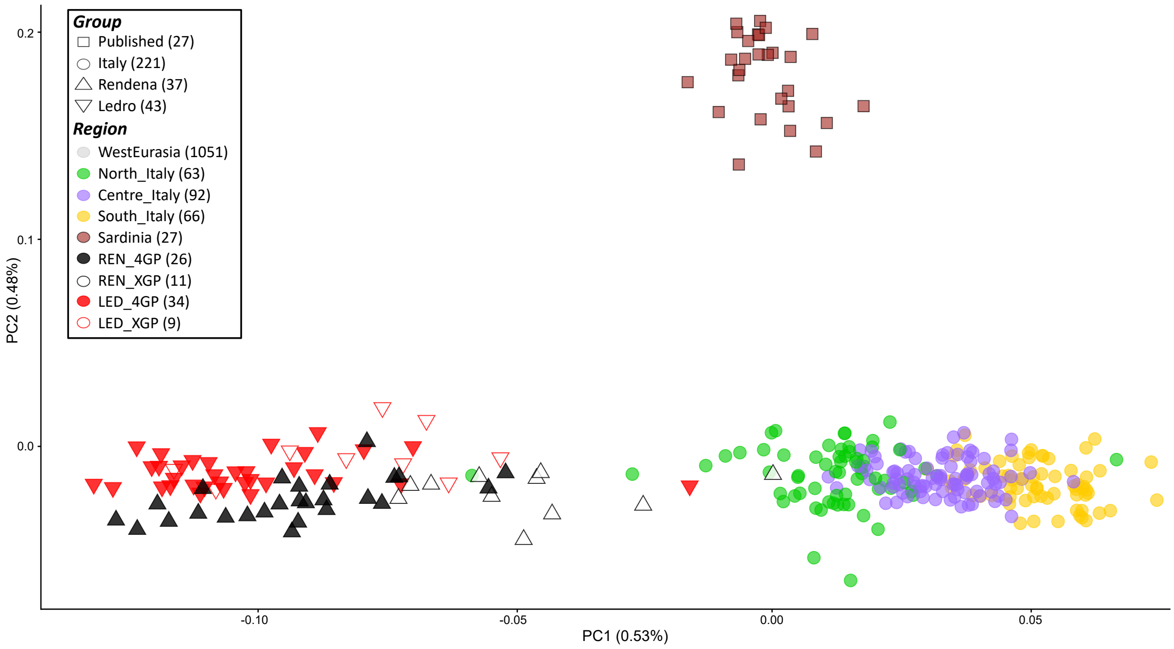


**Fig. S3. PCA of mitochondrial lineage distributions in Italy.** PCA on Italian variability showing PC1 and PC2. Same colours and symbols as main Figures 2–4.

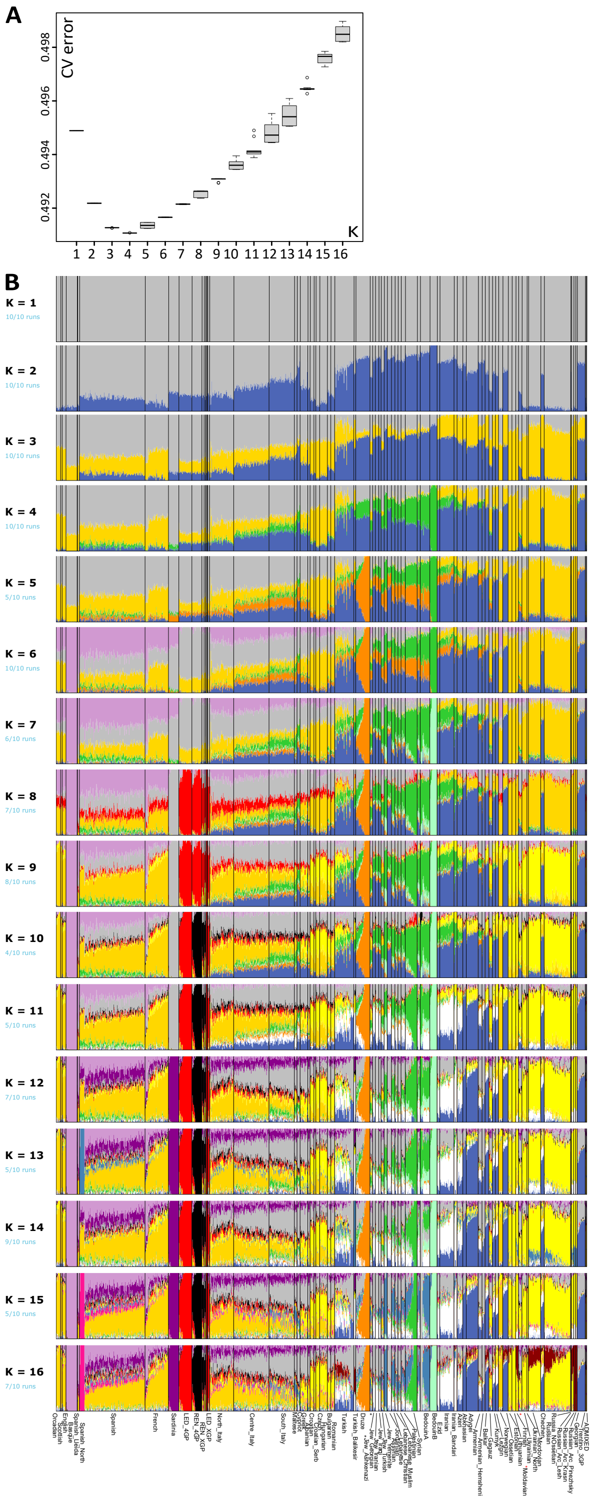

**Fig. S4. Western Eurasian ADMIXTURE.** A) Cross validation error boxplots across Ks. B) ADMIXTURE from K=1 to K=16, 10 replicas each.

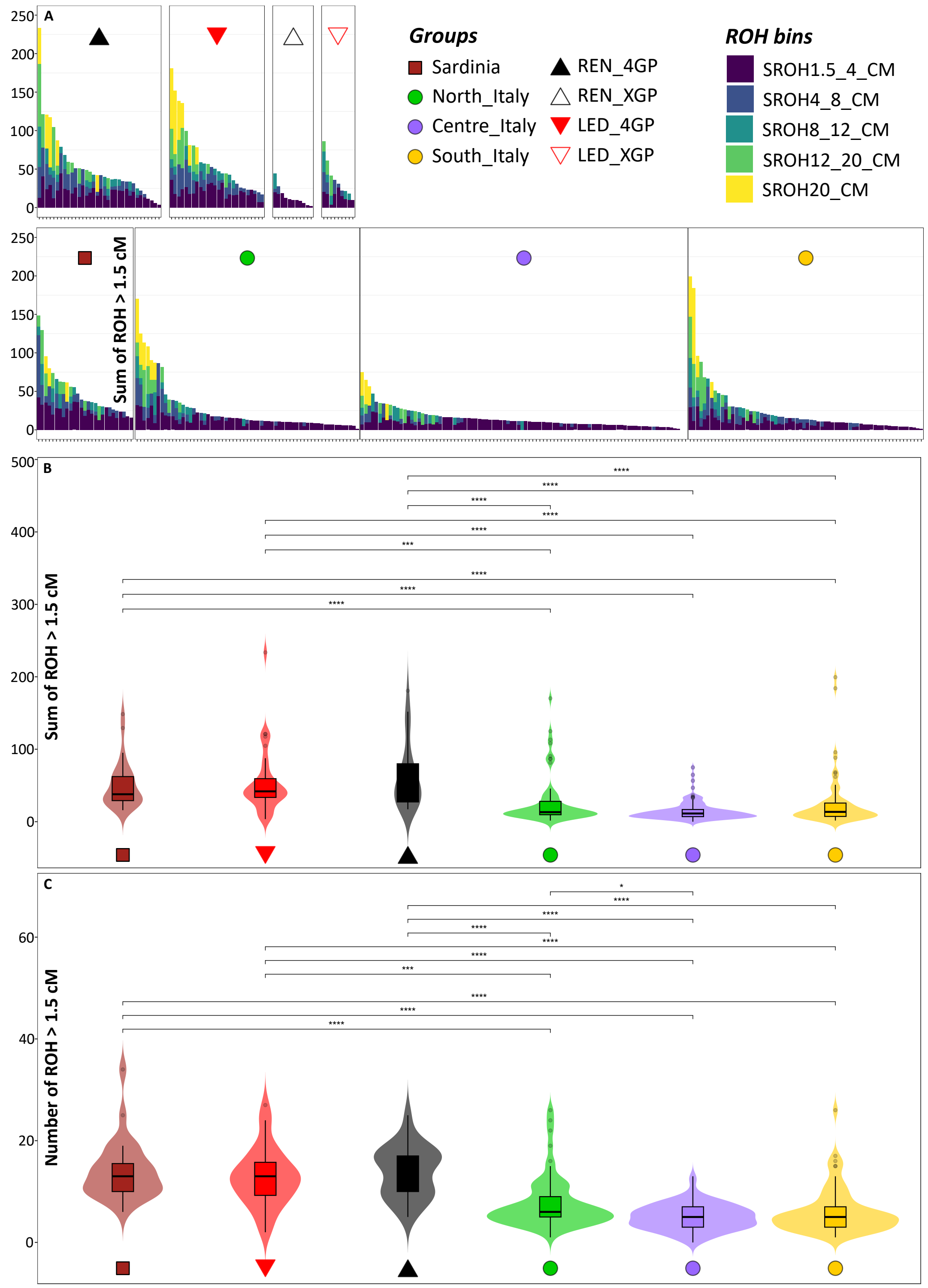


**Fig. S5. ROH.** A) SROH on ROH fragments > 1.5 cM, binned by length, in Italian populations. Legend at top right. B–C) Violin and boxplot distributions of individual-level values of SROH (B) and NROH (C) >1.5 cM across selected Italian populations. Same legend as in A. Pairwise comparisons between populations were assessed using Dunn's test following a

Kruskal–Wallis global test, with Bonferroni correction for multiple testing. Asterisks indicate Bonferroni-adjusted significance levels (* is $p \leq 0.05$, ** $p \leq 0.01$, *** $p \leq 0.001$, **** $p \leq 0.0001$). Only statistically significant pairwise differences (adjusted $p \leq 0.05$) are displayed. Boxes represent interquartile range and median; violin shapes represent kernel density distributions.

A

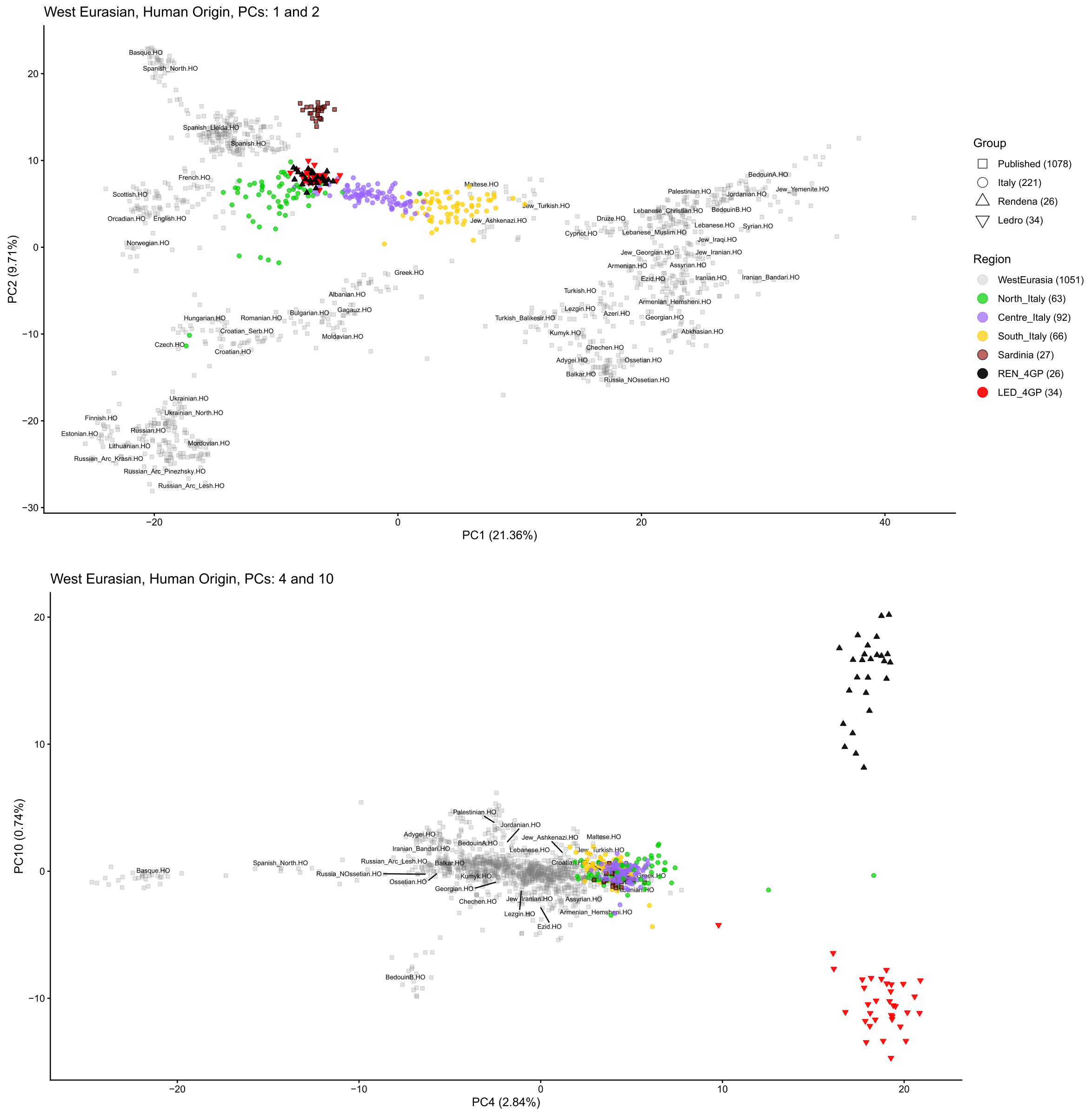


**Fig. S6. PCA on Chromopainter output.** A) PCA showing PC1 and PC2 on ChromoPainter results for Western Eurasia (WE) excluding admixed individuals. Same colours and symbols as main Figures 2–4. Labels for each Western Eurasian population were put at the centroid calculated on that population coordinates. B) PCA showing PC4 and PC10, same legend and labels criteria as in B). PCs were chosen as representatives of several axes of variation that separate Alpine Valleys from WE (PC4) and LED from REN (PC10).

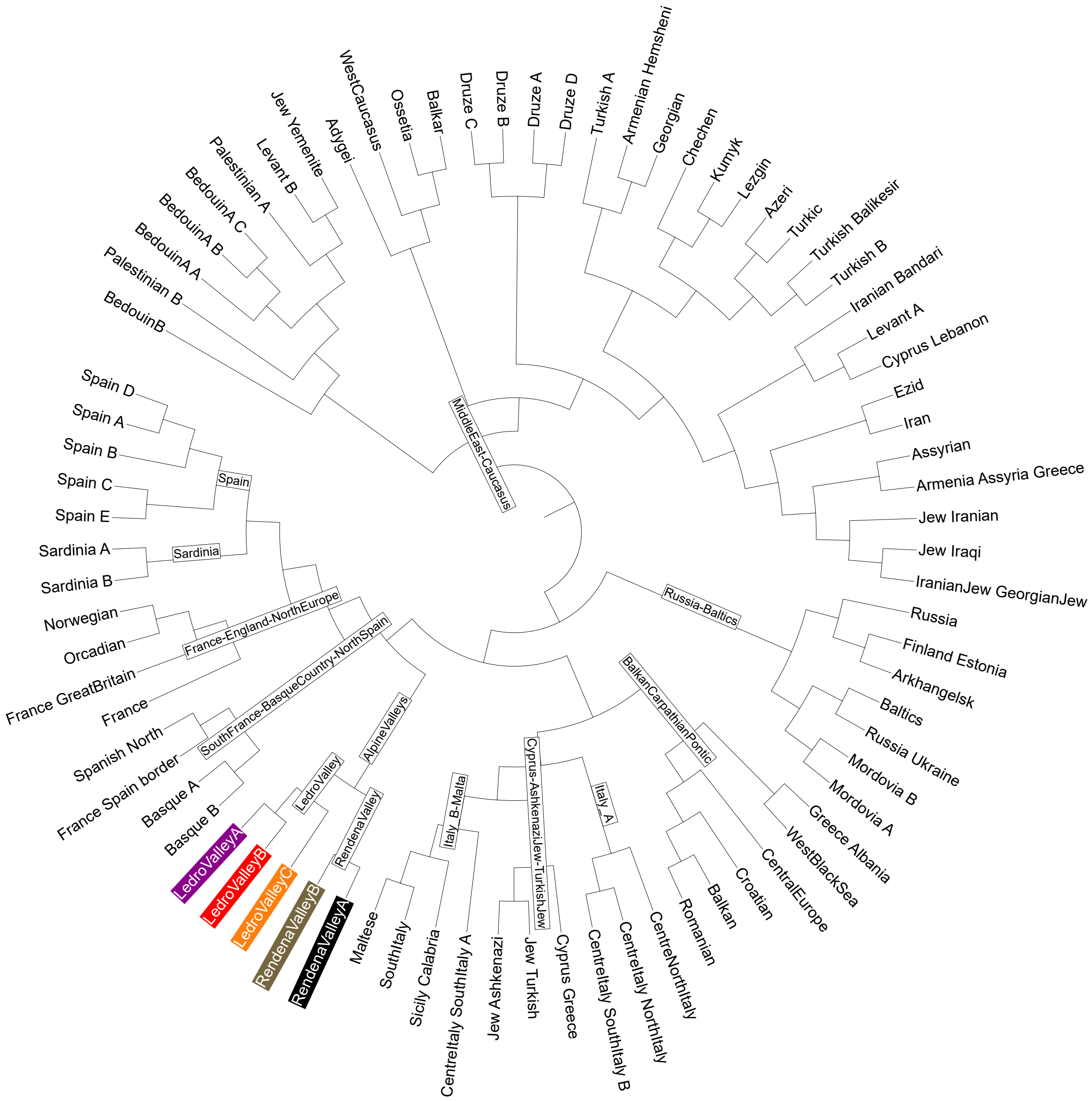


**Fig. S7. Chromopainter/fineSTRUCTURE final cladogram.** Complete cladogram on Western Eurasia as inferred by fineSTRUCTURE algorithm based on ChromoPainter copying vectors. Alpine clusters highlighted. Labels for nodes mentioned or collapsed in main Figure 3A. The tree is unrooted.

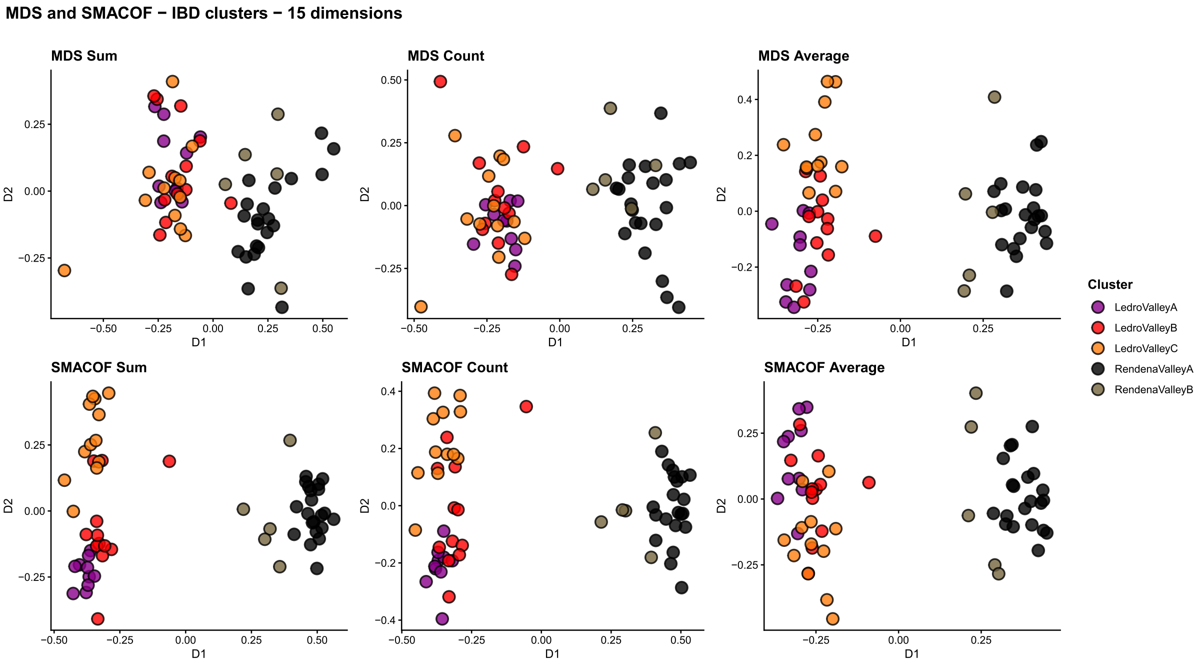


**Fig. S8. Multidimensional scaling (MDS) based on identity-by-descent (IBD) sharing (15 dimensions).** Upper panels: MDS based on the sum, count, and average of shared IBD segments. Lower panels: the same metrics computed using the SMACOF algorithm. The legend is shown on the right.

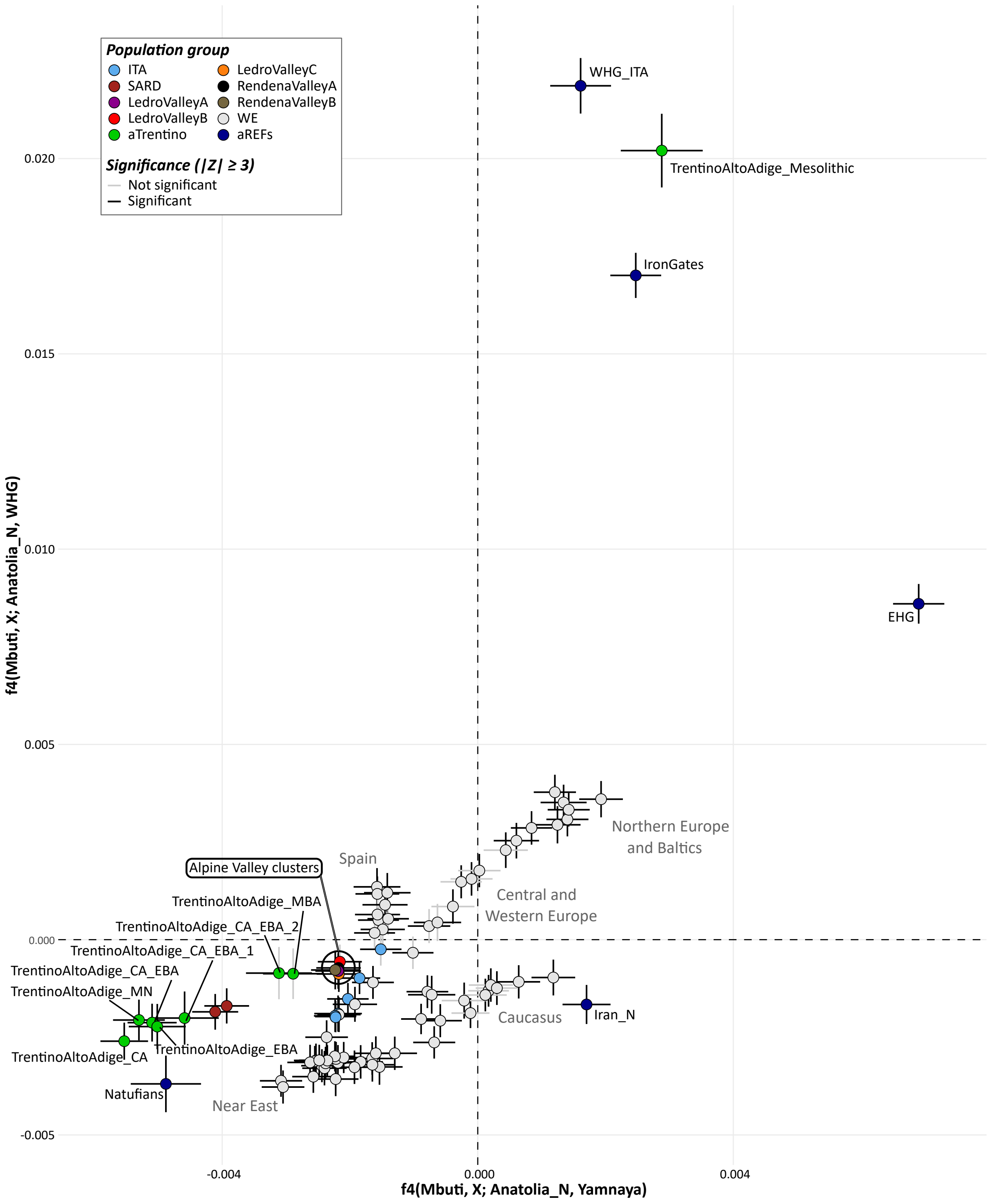


**Fig. S9. Modern Western Eurasia and selected ancient populations affinities.** Affinities toward Anatolia_N relative to either Yamnaya or WHG. Alpine clusters show closest affinity to Italian populations. Error bars represent ±2 standard errors. Legend in inset. aTrentino refers to ancient TAA populations as identified in Croze et al. 2025. aREFs refers to ancient populations that shaped current Western Eurasian genomic variability.

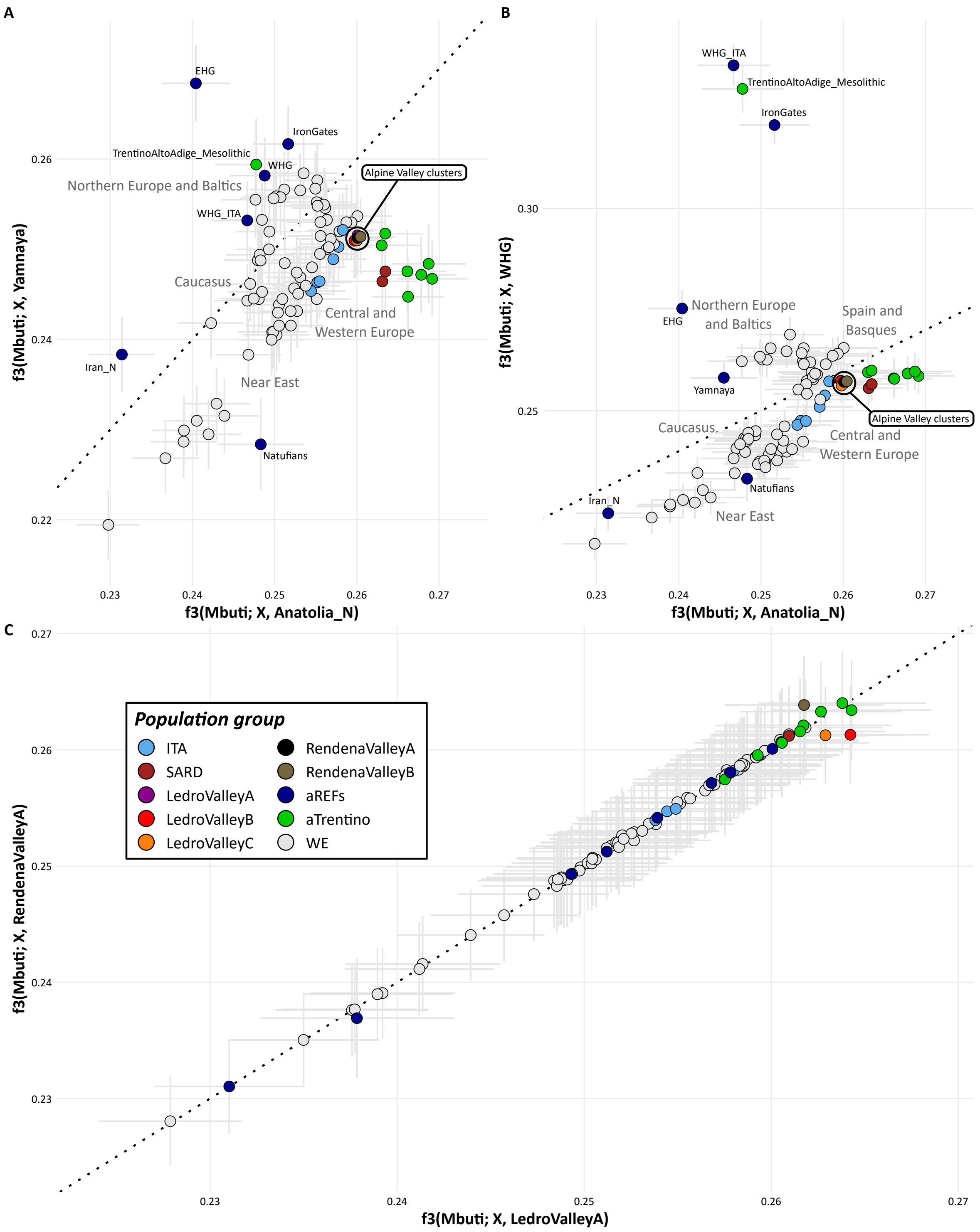


**Fig. S10. Shared drift with f3s.** Biplots with f3(Mbuti; X, Y). X is modern Western Eurasian clusters identified by Chromopainter/fineSTRUCTURE, plus aTrentino (ancient TAA populations as identified in Croze et al. 2025) and aREFs (ancient populations thought to have shaped current Western Eurasian variability). Y is Anatolia_N and Yamnaya (A), Anatolia_N and WHG

(B), LedroValleyA and RendenaValleyA (C). Error bars in light grey represent $\pm 2$ standard errors. Black dotted line is the identity line. Legend in inset of C.
